# Computational modelling of cambium activity provides a regulatory framework for simulating radial plant growth

**DOI:** 10.1101/2020.01.16.908715

**Authors:** Ivan Lebovka, Bruno Hay Mele, Xiaomin Liu, Alexandra Zakieva, Theresa Schlamp, Nial Gursanscky, Roeland M.H. Merks, Ruth Großeholz, Thomas Greb

## Abstract

Precise organization of growing structures is a fundamental process in developmental biology. In plants, radial growth is mediated by the cambium, a stem cell niche continuously producing wood (xylem) and bast (phloem) in a strictly bidirectional manner. While this process contributes large parts to terrestrial biomass, cambium dynamics eludes direct experimental access due to obstacles in live cell imaging. Here, we present a cell-based computational model visualizing cambium activity and integrating the function of central cambium regulators. Performing iterative comparisons of plant and model anatomies, we conclude that the receptor- like kinase PXY and its ligand CLE41 are part of a minimal framework sufficient for instructing tissue organization. By integrating tissue-specific cell wall stability values, we moreover probe the influence of physical constraints on tissue geometry. Our model highlights the role of intercellular communication within the cambium and shows that a limited number of factors is sufficient to create radial growth by bidirectional tissue production.

**Impact statement:** Radial plant growth produces large parts of terrestrial biomass and can be computationally simulated with the help of an instructive framework of intercellular communication loops.

## Introduction

Stem cells in plants are crucial for their longevity and usually maintained in meristems, special cellular environments constituting protective niches [1]. At key positions in the plant body, we find distinct types of meristems that maintain their activity throughout a plant’s life cycle. Shoot and root apical meristems (SAM, RAM) are located at the tips of shoots and roots, respectively, driving longitudinal growth and the formation of primary tissue anatomy in these organs. Moreover, lateral meristems organized in cylindrical domains at the periphery of shoots and roots execute their thickening. The cambium is the most prominent among these lateral meristems [2]. Cambium cells are embedded in two distinct vascular tissues produced in opposite directions by periclinal cell divisions: xylem (wood) and phloem (bast) [3, 4]. These tissues carry out fundamental physiological functions: long- distance transport of water and nutrients in case of the xylem and translocation of sugars and a multitude of signaling molecules in the case of the phloem. Based on its tightly controlled bidirectionality of tissue production and resulting bipartite organization, the cambium is a paradigm for bifacial stem cell niches which produce two tissue types in opposite directions and are found across different kingdoms of life [5].

Balancing proliferation and differentiation within meristems is essential. In the SAM and the RAM this balance is maintained via interaction between the pool of stem cells and the organizing center (OC) and the quiescent center (QC), respectively, where the rate of cell division is relatively low. Both domains form a niche within the meristem instructing surrounding stem cells via regulatory feedback loops [6–10]. In comparison to apical meristems, functional characterization of cambium domains was performed only very recently. During their transition from stem cells to fully differentiated xylem cells, early xylem cells instruct radial patterning of the cambium including stem cell activity and, thus, similar to the OC in the SAM, fulfil this role only transiently [11]. In addition to influence from the early xylem, phloem-derived DNA- BINDING ONE ZINC FINGER (DOF) transcription factors designated as PHLOEM EARLY DOFs (PEARs) move to cambium stem cells and stimulate their proliferation in a non-cell autonomous manner [12]. Furthermore, genetically encoded lineage tracing experiments showed that cell divisions are mostly restricted to individual bifacial stem cells located in the central cambium feeding both xylem and phloem production [11, 13, 14]. Altogether these findings defined functional cambium domains and revealed some of their reciprocal communication.

Another central and well-established mechanism regulating cambium activity in the reference plant *Arabidopsis thaliana* and beyond [15–18] is the action of a receptor- ligand pair formed by the plasma membrane-bound receptor-like kinase PHLOEM INTERCALATED WITH XYLEM (PXY), also known as TDIF RECEPTOR (TDR), and the secreted CLAVATA3/ESR-RELATED 41 (CLE41) and CLE44 peptides. Like the PEAR proteins [12], CLE41 and CLE44 are expressed in the phloem and thought to diffuse to dividing cells in the cambium area expressing PXY [16, 19]. Direct binding of CLE41 to PXY [19–21] promotes the expression of the transcription factor WUSCHEL RELATED HOMEOBOX 4 (WOX4) [22], which, in turn, is crucial for maintaining the capacity of cells to proliferate [15, 22, 23]. At the same time, the PXY/CLE41 module is reported to repress xylem differentiation in a *WOX4*- independent manner [22, 24]. In this context, PXY stimulates the activity of glycogen synthase kinase 3 proteins (GSK3s), like BRASSINOSTEROID-INSENSITIVE 2 (BIN2) [24]. BIN2, in turn, represses the transcriptional regulator BRI1-EMS SUPPRESSOR 1 (BES1), which mediates brassinosteroid (BR) signaling and promotes xylem differentiation [24, 25]. In line with the hypothesis of a dual role in regulating stem cell activity and xylem differentiation, *PXY* is expressed in the proximal cambium zone containing developing xylem cells and in the central cambium zone containing bifacial cambium stem cells [13, 26, 27].

Distally to the *PXY* expression domain and oriented toward the phloem, the closest homolog to PXY, the receptor-like kinase MORE LATERAL GROWTH 1 (MOL1), represses cambium activity [28, 29]. Although their extracellular domains are highly similar, *PXY* and *MOL1* cannot functionally replace each other, indicating that MOL1 activity does not depend on CLE41/44 peptides and that distinct signaling loops act in the proximal and distal cambium domains [29]. The latter conclusion is supported by the finding that the AUXIN RESPONSE FACTOR5 (ARF5) is expressed in the proximal cambium and promotes the transition from stem cells to xylem cells by directly dampening *WOX4* activity [26, 30]. ARF5 activity is enhanced by phosphorylation through the GSK3 BIN2-LIKE 1 (BIL1) which, in contrast to other GSK3s [24], is inhibited by the PXY/CLE41 module [30].

As the role of multiple communication cascades between different cambium-related tissues is beginning to emerge, it is vital to generate a systemic view on their combined impact on cambium activity and patterning integrated into a dynamic tissue environment. However, although the cambium plays an instructive role for stem cell biology, a dynamic view on its activity is missing due to its inaccessibility for live cell imaging. Computational modeling, in particular agent-based modeling combining tissue layout with biochemical signaling processes, can overcome these obstacles and help analyzing the interplay between cellular signaling processes, cell growth and cell differentiation *in silico* that would otherwise be inaccessible. Here, we present a dynamic, agent-based computational model [31] of the cambium integrating the functions of PXY, CLE41, and putative phloem-derived signals into a plant- specific modelling framework. As revealed by informative cambium markers, our model is able to reproduce anatomical features of the cambium in a dynamic manner. It also allows studying the cambium as a flexible system comprised of multiple interacting factors, and the effects of those factors on cell division, differentiation and tissue patterning.

## Results

### Establishing a dynamic cambium model

Taking advantage of the almost exclusive radial expansion of mature plant growth axes, we sought to create a minimal framework recapitulating the 2D dynamics of radial plant growth. To do so, we first produced a simplified stereotypic 2D- representation of a plant growth axis displaying a secondary anatomy by employing VirtualLeaf – a framework specially designed for agent-based modeling of plant tissue growth [32, 33]. To avoid confusion, we refer to factors within the model by an asterisk: e.g., GENE – refers to the plant gene, whereas GENE*\** refers to its model counterpart. Within the model we defined three cell types: Cells designated as cambium*, cells present in the center referred to as xylem*, and cells present distally to the cambium* designated as phloem* (Figure 1A). We then defined rules determining cell* behavior: **i)** all cells* grew until they reached a size specific for each cell type, **ii)** cambium cells* divided when they exceeded a certain size, **iii)** cambium cells* changed their identity into xylem* or phloem* depending on the conditions described below (see also supporting information, Supplementary File 1). All chemical-like factors* implemented in the model had manually chosen cell* type- specific production and degradation rates.

**Figure 1.**
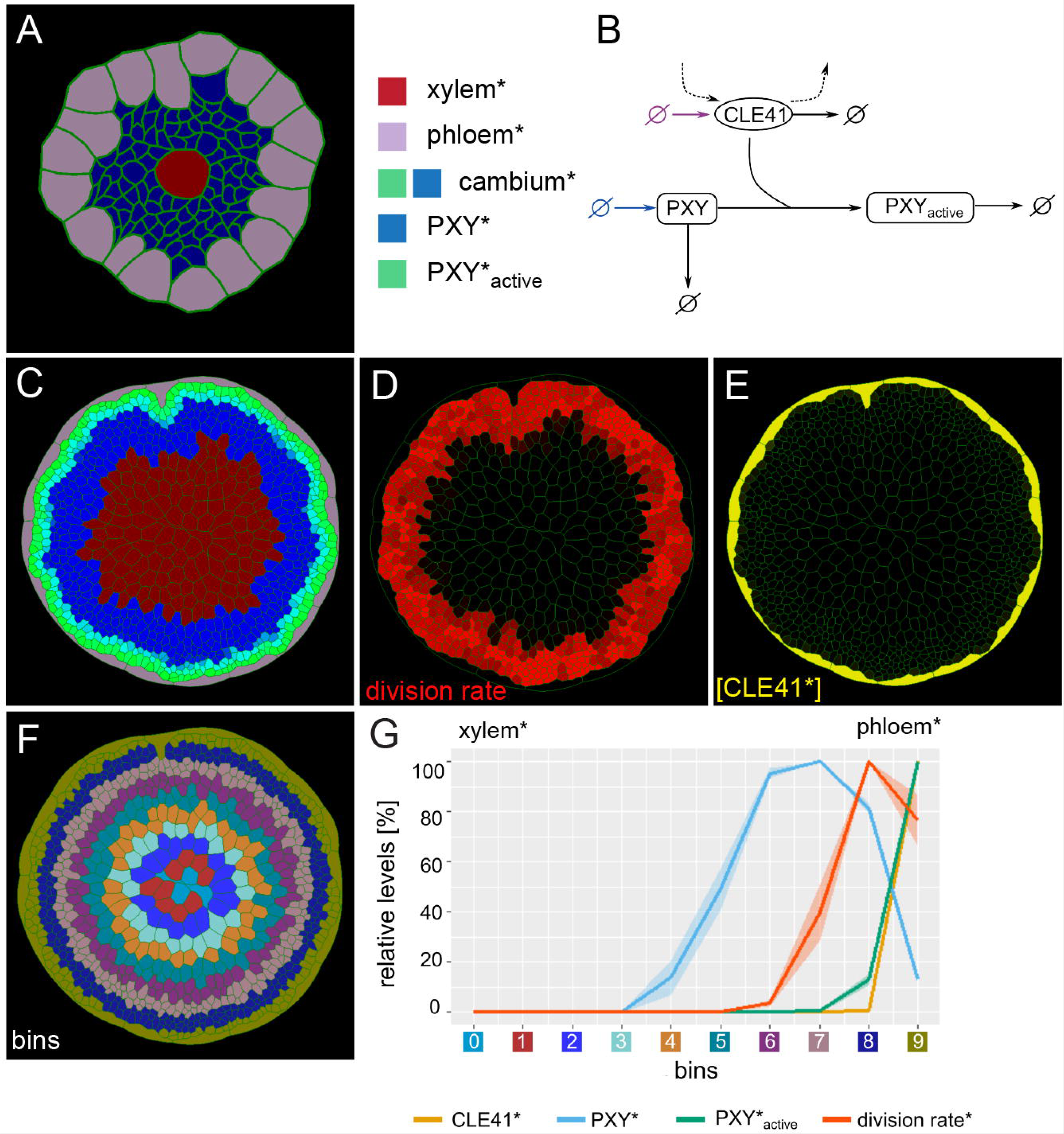
Generation of the initial model. **(A)** Tissue template used to run VirtualLeaf simulations. Phloem* is depicted in purple, xylem* in red. Cambium cells* are colored according to their levels of PXY* and PXY-active*. Cambium* is colored in blue due to the initial level of PXY*. Color legend on the right apply to A and C. **(B)** Schematic representation of the biochemical model. Reactions that occur in all cell types* are drawn in black. Reactions only occurring in the phloem* are depicted in purple, reactions specific to the cambium* are in blue. Crossed circles represent production or degradation of molecules. **(C)** Output of simulation using Model 1. **(D)** Visualization of cell division rates* within the output shown in (C). Dividing cells* are marked by red color fading over time. **(E)** Visualization of CLE41* levels within the output shown in (C). **(F)** Sorting cells* within the output shown in (C) into bins based on how far their centers are from the center of the hypocotyl*. Different colors represent different bins. **(G)** Visualization of the relative chemical levels and division rates in different bins shown in (F) averaged over n=10 simulations of Model 1. Each chemical’s bin average is calculated and then expressed as a percentage of the maximum bin value of the chemical. Bin colors along the x-axis correspond to the colors of bins in (F).

To implement context-dependent regulation of cambial cell division and differentiation, we took advantage of the PXY/CLE41 signaling module [19, 22]: Phloem cells* produced a factor designated as CLE41* able to diffuse between cells*, whereas the corresponding, non-diffusing receptor designated as PXY* is produced in cambium cells* (Figure 1B). Recapitulating the CLE41-dependent function of PXY, we considered the following reaction:

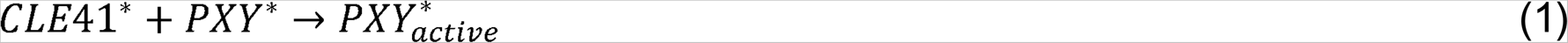

Thereby, the presence of both CLE41* and PXY* in a cell turned PXY* into PXY_active_* (Figure 1B). For cambium cells* we described the PXY*-CLE41* interaction by the following equations:

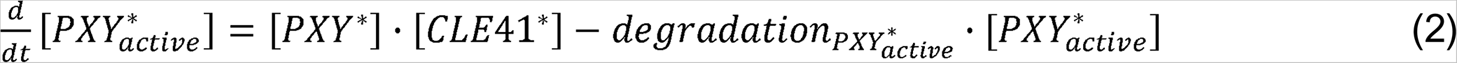

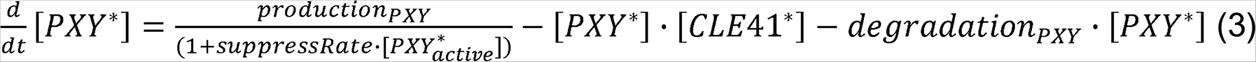

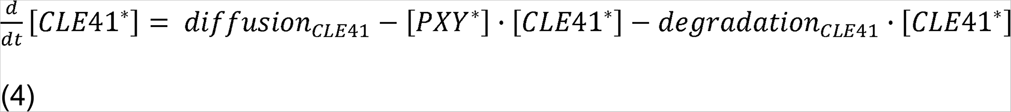

In these equations, [X*] denoted the concentration of the respective factor in each cell*. Since PXY-CLE41 signaling was reported to negatively regulate *PXY* expression [16], we assumed that the production rate of PXY* is inhibited by [PXY_active_*]. Therefore, the higher [PXY_active_*] in a given cell*, the less PXY* was produced (equation 3). To integrate PXY/CLE41-dependent regulation of cell proliferation, we let cambium cells* divide only when [PXY_active_*] exceeded a certain threshold. Thereby, the proliferation of cambium cells* was dependent on both, locally produced PXY* and CLE41* originating from the phloem*. To instruct the differentiation of cambium cells*, we took advantage of the observation that the PXY/CLE41 module represses xylem differentiation [19, 24]. Consequently, we instructed cambium cells* to change their identity into xylem* as soon as they reached a certain size and [PXY_active_*] became lower than a threshold value (Supplementary file 1).

In the resulting Model 1, the growing structure maintained a circular pool of dividing cambium cells* with a high concentration of PXY_active_* while producing xylem cells* toward the center of the organ (Figure 1C, Video 1, Video 2, Video 3, Video 4, Supplementary Information Items). As expected, when cambium cells* were displaced to the proximal side of the cambium*, they stopped dividing likely due to low [PXY_active_*] (Figure 1C, D, Video 3, Video 4, Video 5) allowing them to reach a size sufficient for xylem* differentiation. Cell* division rates were highest close to CLE41* producing phloem cells* (Figure 1D-G, Video 2, Video 3). Moreover, as PXY_active_* negatively affected the production of PXY*, [PXY*] was particularly low in the distal cambium* region (Figure 1C, F, G. Video 1, Video 5). This pattern was reminiscent of the exclusive activity of the *PXY* promoter in the proximal cambium area observed previously [13, 29]. Thus, although phloem was not produced, with maintaining a circular domain of cambium cells* and cell* proliferation and with promoting xylem* production, Model 1 was able to recapitulate several core features of the active cambium.

### The combination of *PXY* and *SMXL5* promoter reporters reveals cambium anatomy

To identify rules for phloem formation, we took advantage of findings obtained using the *PXYpro:CYAN FLUORESCENT PROTEIN* (*PXYpro:CFP*) and *SUPPRESSOR OF MAX2-LIKE 5pro:YELLOW FLUORESCENT PROTEIN* (*SMLX5pro:YFP*) markers, recently established read-outs for cambium anatomy [13]. *PXYpro:CFP* and *SMXL5pro:YFP* markers label the proximal and distal cambium domain (Figure 2A), respectively, and are therefore indicative of a bipartite cambium organization. *PXYpro:CFP* activity indicates the proximal xylem formation zone whereas *SMXL5pro:YFP* activity indicates the distal phloem formation zone. A narrow central zone in which both markers are active hold cambium stem cells which feed both tissues and also show a high rate of cell divisions in comparison to xylem and phloem progenitors [13].

**Figure 2.**
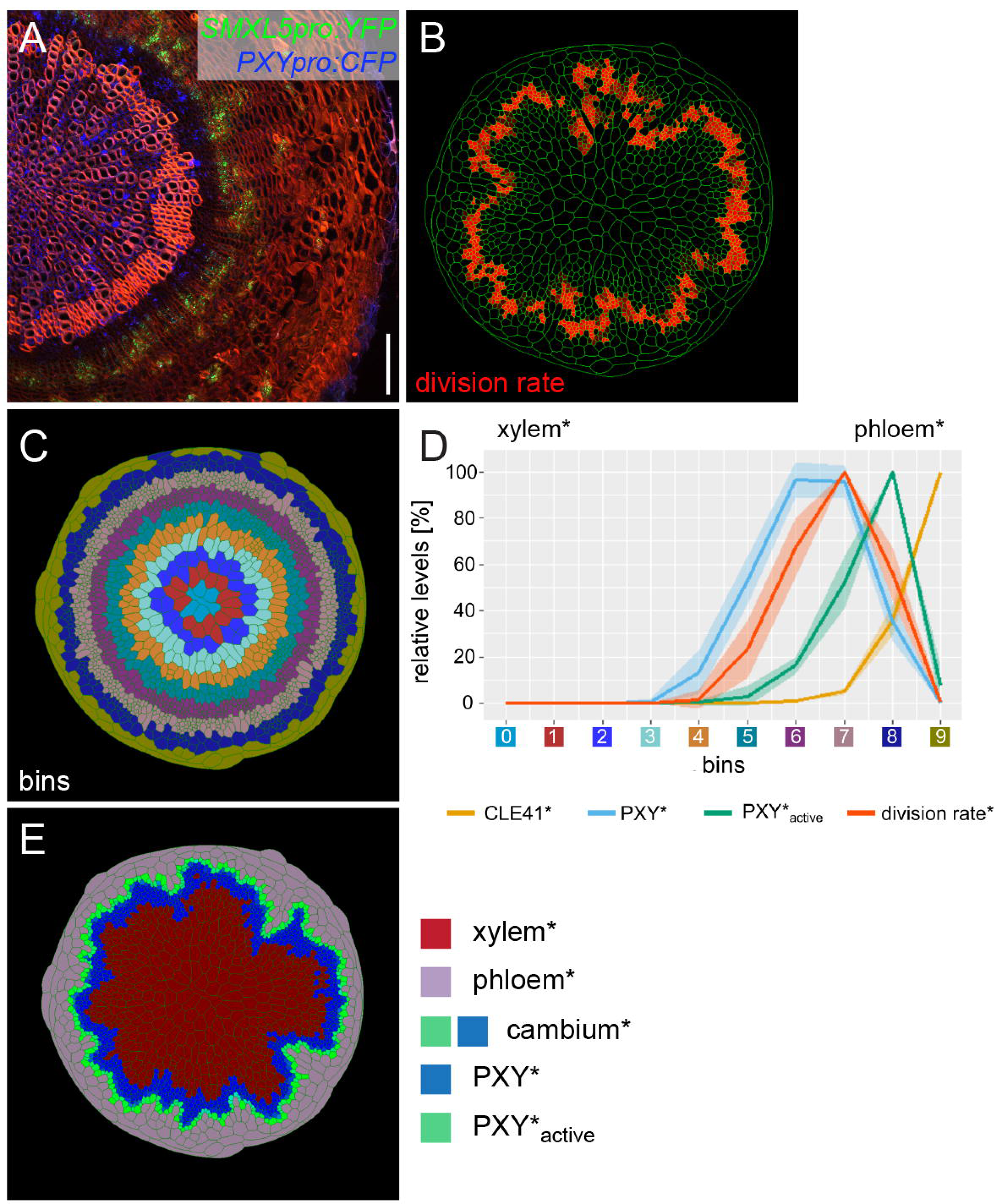
Implementing phloem formation into the model. **(A)** Cross-section of a wild type hypocotyl expressing *PXYpro:CFP* (blue) and *SMXL5pro:YFP* (green). Cell walls are stained by Direct Red 23, mainly visualizing xylem (red). Only a quarter of the hypocotyl is shown with the center in the bottom right corner. Scale bar: 100 µm. **(B)** Visualization of cell division rates* within the output shown in (C and E). Dividing cells* are marked by red color fading over time. **(C)** Sorting cells* within the output shown in (B and E) into bins. **(D)** Visualization of the average relative chemical levels* and division rates* in different bins of repeated simulations of Model 2A (n=10). Bin label colors along the x-axis correspond to the colors of bins shown in (C). **(E)** Output of simulation using Model 2A. Unlike Model 1, Model 2A produces new phloem cells*.

To computationally recapitulate the observed maximum of cell division rates in the central cambium domain, we sought to inhibit cell* divisions in the distal layers of the cambium*. Such an effect is, for instance, mediated by the receptor-like kinase MOL1, which, similarly to *SMXL5*, is expressed distally to *PXY* expressing cells and suppresses cambial cell divisions [29]. Because cells* in the distal cambium* region were characterized by high levels of PXY_active_* (Figure 1C, G, Video 4), we used PXY_active_* to locally inhibit cell* division and, at the same time, to instruct phloem* formation. Therefore, we modified the rule for cell* differentiation such that, when a cambium cell* reached a specific size, it differentiated into xylem* if [PXY_active_*] became lower than a threshold value and into phloem* if [PXY_active_*] was greater than the same threshold and the cell was larger (Supplementary File 1). Thereby, our model followed a classical ‘French flag’ principle of development according to which concentration gradients of diffusible morphogens pattern surrounding tissues [34]. It is worth noting that the combined effect of CLE41* on cell* proliferation, on phloem* specification and on [PXY*] may also be achieved by distinct phloem-derived factors mediating these effects individually.

The computational implementation of these rules (Model 2A) resulted in a descending gradient of cell* division rates in the distal cambium* domain likely due to high levels of PXY_active_* (Figure 2B-D, Video 6, Video 7, Video 8). The cell* division rate was highest in the central cambium* domain defined by high [PXY*] and by moderate [PXY_active_*] (Figure 2B-D, Video 8, Video 9, Video 10, Video 11). Also, not only xylem* but also phloem* was continuously produced and the fate of cambium cells* was dependent on their position relative to the differentiated tissues*. In the central cambium* domain, cells* proliferated and constantly replenished the stem cell pool (Figure 2B, Video 6, Video 7, Video 8). Thus, by incorporating relatively simple rules, Model 2A was able to recapitulate major cambium features, including phloem formation. Moreover, in qualitative terms, the resulting anatomy* reproduced the anatomy of a mature *Arabidopsis* hypocotyl (Figure 2A, E).

### Cambium model explains the effect of ectopic CLE41 expression

To evaluate the predictive power of Model 2, we tested its capacity to simulate the effects of experimental perturbation of cambium regulation. Ectopic expression of *CLE41* by employing the *IRREGULAR XYLEM 3/CELLULOSE SYNTHASE CATALYTIC SUBUNIT 7* (*IRX3/CESA7*) promoter, which is active in cells undergoing secondary cell wall deposition [35–37], substantially alters hypocotyl anatomy [16]. This effect was confirmed when *PXYpro:CFP/SMXL5pro:YFP* activities were analyzed in a plant line carrying also an *IRX3pro:CLE41* transgene (Figure 3A). The *PXYpro:CFP* activity domain had a cylindrical shape including the proximal cambium domain and the xylem tissue itself in plants with a wild type background (Figure 2A).

**Figure 3.**
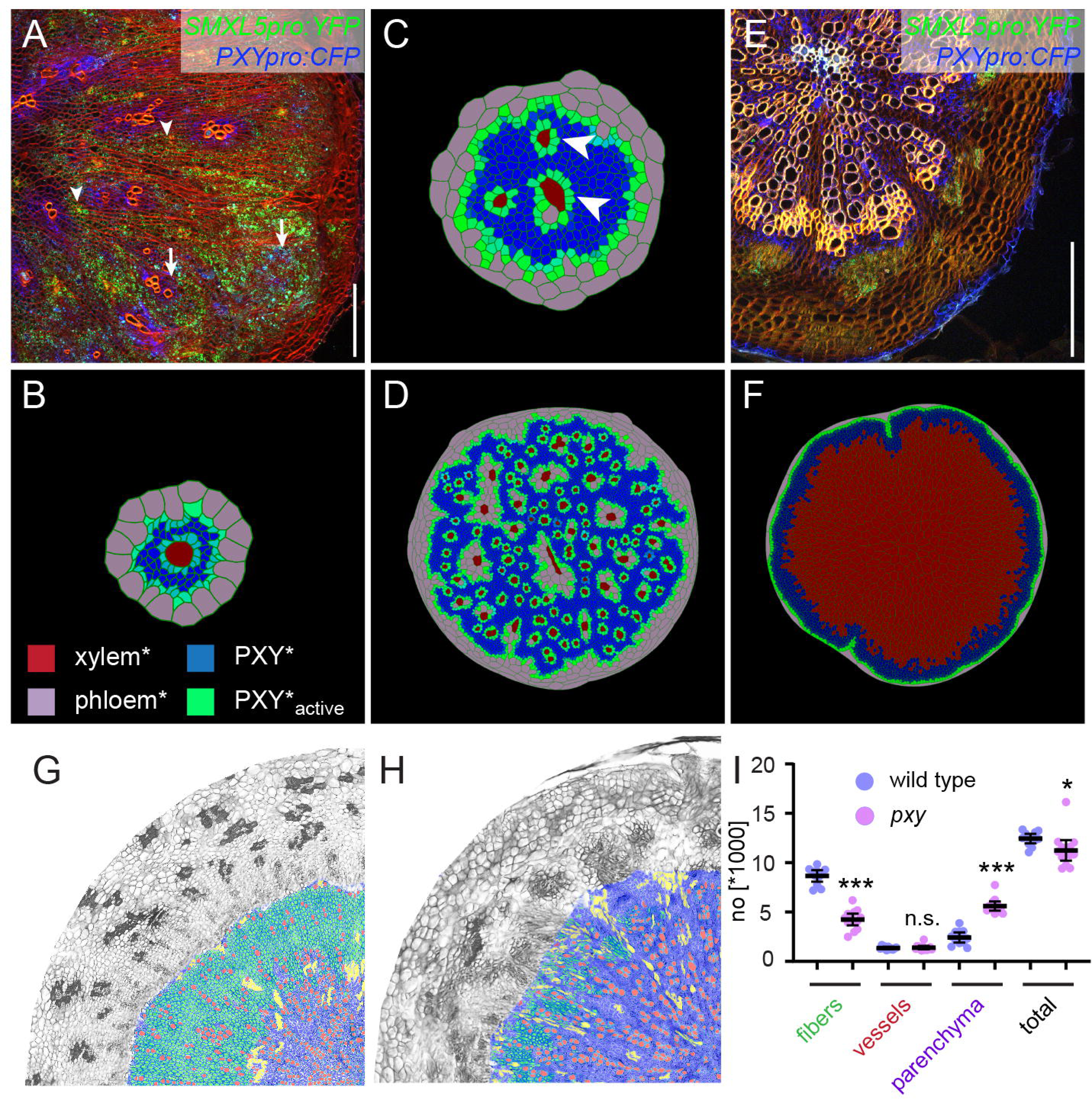
Comparing the effect of perturbing cambium activity in the model and in plants. **(A)** Cross-section of a hypocotyl carrying *PXYpro:CFP* (blue), *SMXL5pro:YFP* (green) markers, and the *IRX3pro:CLE41* transgene. Cell walls are stained by Direct Red 23 visualizing mostly xylem (red). Arrowheads point to proximal hypocotyl regions where *SMXL5pro:YFP* activity is found. Arrows indicate distal regions with *SMXL5pro:YFP* activity. Cell walls are stained by Direct Red 23 visualizing mostly xylem (red). Only a quarter of the hypocotyl is shown with the center in the upper left corner. Scale bar: 100 µm. **(B)** First frames of Model 2B simulations. Due to the expression of CLE41* by xylem cells*, high levels of PXY-active* are generated around xylem cells* already at this early stage. Legend in B indicates color code in B, C, D, F. **(C)** Intermediate frames of Model 2B simulations. Newly formed xylem cells* express CLE41* and produce high levels of PXY-active* next to them (white arrowheads). **(D)** The final result of Model 2B simulations. Zones of PXY* (blue) and PXY- active*(green) are intermixed, xylem cells* are scattered, and phloem cells* are present in proximal areas of the hypocotyl*. **(E)** Cross-section of a *pxy* mutant hypocotyl carrying *PXYpro:CFP* (blue) and *SMXL5pro:YFP* (green) markers, stained by Direct Red 23 (red). The xylem shows a ray-like structure. Only a quarter of the hypocotyl is shown with the center in the upper left corner. Scale bar: 100 µm. **(F)** Final result of Model 2D simulations. Reducing PXY* levels leads to similar results as produced by Model 1 (Figure 1B) where only xylem* is produced. **(G, H)** Comparison of histological cross-sections of a wild type (G) and a *pxy* (H) mutant hypocotyl, including cell type classification produced by ilastik. The ilastik classifier module was trained to identify xylem vessels (red), fibers (green), and parenchyma (purple), unclassified objects are shown in yellow. **(I)** Comparison of the number of xylem vessels, fibers and parenchyma cells found in wild type (blue) and *pxy* mutants (purple). Welch’s t test was performed comparing wild-type and *pxy* mutants for the different cell types (n = 11-13). \*\*\**p* < 0.0001, \**p* < 0.05. Lines indicate means with a 95 % confidence interval. 11-12 plants each for wild type and *pxy* mutants were compared.

While in the presence of the *IRX3pro:CLE41* transgene, *PXYpro:CFP* activity marked a reduced overall number of differentiated xylem cells and was present in irregularly shaped patches distributed over the whole cross-section (Figure 3A). Moreover, we observed regions without *PXYpro:CFP* activity in proximal hypocotyl regions where *SMXL5pro:YFP* was active (Figure 3A). Besides, a substantial part of *SMXL5pro:YFP* activity was detected in the distal regions of the hypocotyl forming islands of irregular shape sometimes intermingled with *PXYpro:CFP* activity (Figure 3A). This activity pattern was in contrast to the one found in plants without the *IRX3pro:CLE41* transgene where *SMXL5pro:YFP* reporter activity surrounded the *PXYpro:CFP* expression domain only from the distal side (Figure 2A). These results indicated that not only the radial symmetry of the hypocotyl [16] but also cambium organization depends on the site of CLE41 production.

To simulate the effect of the *IRX3pro:CLE41* transgene *in silico*, we instructed xylem* cells* to produce CLE41* at the same rate as phloem cells* (Model 2B). Although in this case xylem* formation was initially repressed possibly due to high levels of PXY_active_* in all cambium* cells (Figure 3B, Video 12, Video 13, Video 14), new xylem cells* were formed as soon as the distance between existing xylem and phloem cells* became large enough such that CLE41* levels and, in turn, [PXY_active_*] dropped to permissive levels (Figure 3C, Video 12, Video 13, Video 14). New phloem cells* were produced close to existing phloem and xylem cells* likely due to high levels of PXY_active_* (Figure 3C, Video 15, Video 16). As a result, Model 2B produced a similar disruption in cambium* organization, as observed in *IRX3pro:CLE41* plants (Figure 3D, Video 12, Video 13, Video 14). Zones with both high [PXY_active_*] and low [PXY*], which were found in the distal cambium* in Model 2A (Figure 2B, Video 16, Video 17), appeared in the organ* center together with individual xylem cells* (Figure 3D).

Moreover, in addition to being produced in distal regions, new phloem cells* were produced in the central areas of the organ* as demonstrated previously for *IRX3pro:CLE41* plants [16]. Thus, rules determining cambium* polarity implemented in Model 2 were sufficient to simulate organ anatomy found in wild type and *IRX3pro:CLE41* genetic backgrounds.

In contrast, a discrepancy between the model logic and the *in planta* situation was suggested when we compared a model having reduced PXY* activity with *pxy* mutants carrying the *PXYpro:CFP* and *SMXL5pro:YFP* reporters. In *pxy* mutants, the xylem tissue did not have a cylindrical shape, but was instead clustered in radial sectors showing *PXYpro:CFP* and *SMXL5pro:YFP* activity at their distal ends, whereas regions in between those sectors had little to no xylem and no (Figure 3E). *PXY* promoter reporter activity was observed distally to xylem sectors, whereas the *SMXL5* promoter activity was as usual present distally to the *PXY* activity domain. Interestingly, *PXYpro:CFP* and *SMXL5pro:YFP* activity domains were still completely distinct meaning that *PXYpro:CFP* activity did not expand into the *SMXL5pro:YFP* domain (Figure 3E). This discrepancy indicated that, in contrast to our assumption, the CLE41-PXY signaling module did not restrict *PXY* promoter activity in the distal cambium. The discrepancy between Model 2 and the situation in plants was confirmed when we completely eliminated PXY* activity from our model (Model 2C). As expected, this elimination resulted in the absence of growth due to the full dependence of cell* divisions on the PXY* function, clearly being at odds with the phenotype of *pxy* mutants (Figure 3E). Even when we only reduced PXY* activity (Model 2D), this did not result in a split of the continuous cambium domain* but abolished phloem* formation and increased the production of xylem (Figure 3F).

Interestingly, the quantification of water transporting xylem vessels, xylem fibers, which provide mechanical stability, and xylem parenchyma in sections from wild type and *pxy* mutant hypocotyls by automated image segmentation revealed that the total number of xylem cells and the number of xylem vessels was comparable (Figure 3G- I, Figure 3-figure supplement 1). In contrast, the number of cells classified as fibers was substantially reduced in *pxy* mutants whereas the number of cells classified as parenchyma was increased (Figure 3G-I). These results suggested that during radial growth, *PXY* promotes the formation of xylem fibers, while the formation of xylem vessels and the total number of cambium-derived cells produced toward the xylem is hardly *PXY*-dependent.

### Multiple phloem-derived factors determine cambium activity

Our observations prompted us to reconsider some features of the model and to extend our ‘French flag’ approach. As the proximal cell production rate by the cambium was not *PXY*-dependent, we made xylem* formation independent from the control of PXY-active*. Instead, cambium cells* differentiated into xylem cells* when they reached a specific size and, at the same time, expressed PXY* as a positional feature. To maintain a population of active cambium cells* in the absence of PXY*, we introduced a second phloem*-derived factor (PF), reminiscent of the PEAR transcription factors identified recently [12]. PF* stimulated cell* divisions by promoting the production of a division factor (DF) in cambium cells* and in phloem parenchyma* (Figure 4A, see below). Cambium cells* divided only if the concentration of DF* exceeded a threshold value (Supplementary File 1). DF* production was at the same time stimulated by PXY_active_* as its only effect in cambium cells* (Figure 4A). Thereby, cambial cell* divisions were dependent on the combined influence of PXY_active_* and their proximity to phloem poles* (see below).

**Figure 4.**
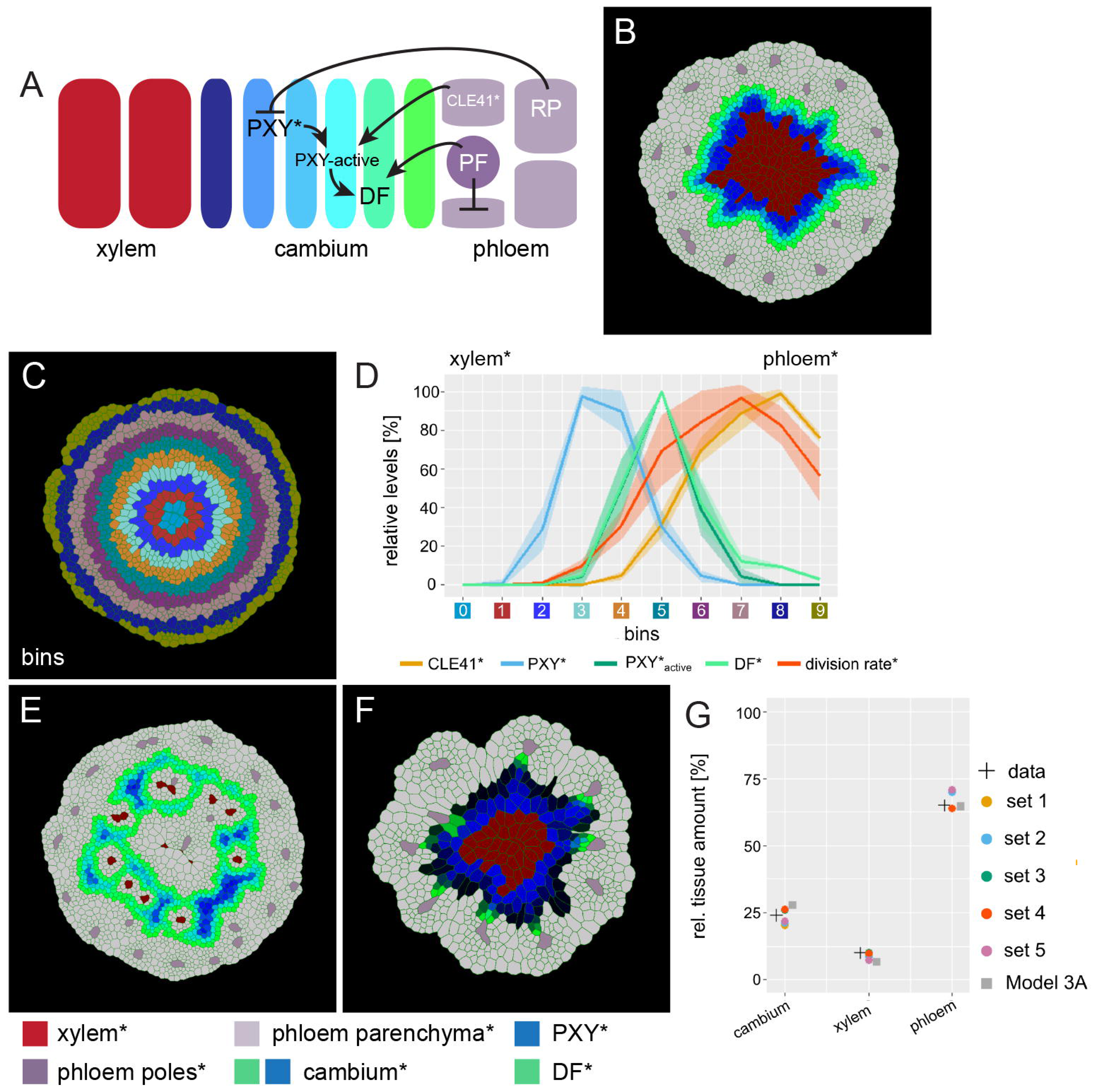
An extended model for simulating genetic perturbations. **(A)** Regulatory network proposed based on experimental observations. **(B)** Result of the simulation run for Model 3A. This model implements the network interactions described in (A). Color coding at the bottom of Figure 4. **(C)** Outline of cell bins for the results of Model 3A, as shown in (B). **(D)** Visualization of the relative levels of chemicals* and division rates* in different bins. Bin colors along the x-axis correspond to the bin colors in (C). **(E)** Output of Model 3B simulation. Ectopic CLE41* expression was achieved by letting xylem cells* produce CLE41*. **(F)** Output of Model 3C. Simulation of the *pxy* mutant was achieved by removing the stimulation of DF* production by PXY* and hence by removing the effect of PXY* on cell division and cambium* subdomain patterning. Because of the network structure, PXY* can be eliminated from Model 3 without letting the model collapse (Figure 4F) but reproducing the *pxy* mutant phenotype observed in adult hypocotyls (Figure 4E). **(G)** Estimated tissue ratios for five identified parameter sets compared to experimental values (‘data’) found for wild type (Col-0) hypocotyls 20 days after germination [38] and compared to the final model output before the automated parameter search (’Model 3A’) and the implementation of experimentally determined cell wall thickness for xylem* and phloem*.

PF* was, thus, produced in phloem poles* and the levels in other cells* were determined by the diffusion and degradation:

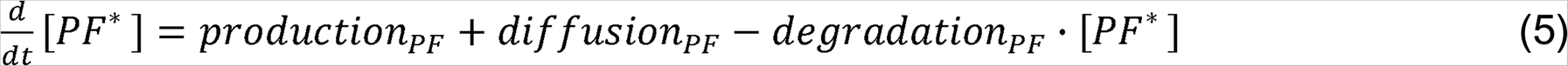

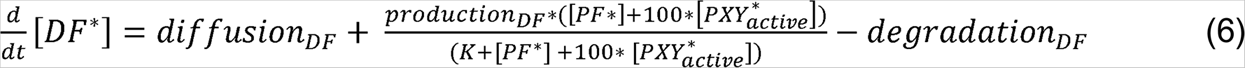

Where K stands for an empirically defined parameter capping the production rate of DF*.

Based on the strong association of xylem sectors with developing phloem cells (Figure 3E), we further hypothesized that the formation of those sectors in *pxy* mutants was dependent on the heterogeneity of cell type distribution in the phloem. Therefore, phloem cells* from the previous models were split into two cell types – phloem parenchyma* and phloem poles* (Figure 4A). To achieve the dispersed pattern of phloem poles, cambium-derived cells* fulfilling the criteria to differentiate into phloem* (see above), differentiated into phloem poles* by default, unless inhibited by PF*, which was specifically produced in pole cells* (Supplementary File 1). Thereby, phloem poles* suppressed phloem pole* formation in their vicinity, expected to result in a patchy pattern of phloem poles as observed *in planta* [38]. The inhibition of phloem poles in their immediate environment is reminiscent to the CLE45/RECEPTOR LIKE PROTEIN KINASE 2 (RPK2) signaling cascade restricting protophloem sieve element identity to its usual position [39, 40]. It is worth noting that in our model, CLE41* was still produced in both phloem poles* and phloem parenchyma* but with a higher rate in phloem poles*. To further achieve PXY*- independent cambium subdomain separation, phloem parenchyma* and phloem poles* were set to express another diffusive signal (RP) which suppressed PXY* expression in cambium* cells, the role that was played by PXY_active_* before (Figure 4A, Supplementary File 1). The role of RP is reminiscent to the role of cytokinin which inhibits xylem-related features in tissue domains designated for phloem development [41]. Importantly, cell divisions in the distal cambium* were not actively repressed anymore but were exclusively dependent on cell size and the level of DF* (Supplementary File 1).

The implementation of these principles *in silico* (Model 3A) resulted again in the establishment of two cambium* subdomains – the distal subdomain which was characterized by high concentrations of DF* and the proximal subdomain characterized by high PXY* concentration (Figure 4B-D, Video 18, Video 19, Video 20, Video 21, Video 22, Video 23). Distally, the cambium* produced phloem parenchyma cells* from which phloem poles* were continuously formed with a pattern resembling the patchy phloem pattern observed in plants (Figure 4B) [38, 42]. Interestingly, the localization of PF* production mainly in phloem poles* resulted in increased DF levels in the vicinity of those poles and, consequently, in locally increased cell* division rates (Video 20, Video 21). This observation is in line with the observation that phloem poles drive cell divisions in their immediate environment and that phloem cells still divide after initial specification [12, 14]. When comparing the radial pattern of *PXYpro:CFP/SMXL5pro:YFP* activities and, as an *in silico* approximation to these activities, the distribution of PXY* and DF* in our model over time, patterns were stable in both cases (Figure 4B, Figure 3-figure supplement 2, Video 18, Video 22, Video 23). This demonstrated that our model was able to generate stable radial patterns of gene* activity comparable to the *in planta* situation.

By instructing CLE41* production additionally in xylem cells*, we next simulated CLE41-misexpression by the *IRX3* promoter (Model 3B, Figure 4E, Video 24, Video 25, Video 26, Video 27, Video 28, Video 29). CLE41* interacted with PXY* on the proximal cambium* border, which resulted in ectopic DF* production and phloem- parenchyma* formation in the proximal hypocotyl* regions (Figure 4E, Video 24, Video 28), similarly as during radial hypocotyl growth in *IRX3pro:CLE41* plants (Figure 3A, Figure 3-figure supplement 2). Still, xylem cells* were formed, generating a patchy xylem* pattern resembling the xylem configuration found in *IRX3pro:CLE41* plants (Figure 3A, Figure 4E, Video 24).

Fully eliminating PXY* activity but keeping the positional information of PXY* for xylem cell differentiation (Model 3C) generated a patchy outline of the distal cambium* subdomain (Figure 4F, Video 30, Video 31, Video 32, Video 33, Video 34, Video 35). While PXY* was usually the main trigger of cell* divisions in cambium cells* at a certain distance from phloem poles*, PF* was sufficient for triggering cell divisions next to phloem poles*. Heterogeneous cambium activity was already observable at early phases of radial hypocotyl growth *in silico* and *in planta* and resulted overall in a reduced tissue production in both systems (Figure 4F, Figure 3- figure supplement 2, Video 30, Video 31, Video 32, Video 33, Video 34, Video 35).

Thus, by introducing both a PXY*-independent pathway stimulating cambium* proliferation and a dependence of cell* proliferation on the distance to phloem poles*, we were able to simulate important features of the *pxy* mutant phenotype (Figure 3E, Figure 4F, Figure 3-figure supplement 2). Collectively, we concluded that we established a computational cambium model sufficiently robust to simulate major genetic perturbations of cambium regulation.

### Physical properties of cambium-derived cells have the potential to influence stem cell behavior

Next, we were interested to see whether the established model was able to reveal organ-wide features of radial plant growth. A characteristic of cambium stem cells is that they divide mostly in periclinal orientation, which is in parallel to the organ surface, resulting in the frequent formation of radial cell files (Figure 2A). Interestingly, although the overall tissue anatomy of the modelled organ* resembled the *in planta* situation, cell division orientation in our model outputs was almost random suggesting that radial cell file formation cannot be explained by the molecular signaling pathways implemented into the model (Figure 4A). The strong dominance of periclinal divisions *in planta*, however, implies the presence of a positional signal instructing cell division orientation. Because classical observations indicated that physical forces play a role in this regard [43–45], we tested whether the model was suited for finding indications for the influence of differential cell stiffness on geometric features of radial plant growth.

To do so, we first determined the relative cell wall thickness in hypocotyl cross sections using the cell wall dye Direct Red 23 [46] as an indication. Notably, staining intensity and, thus, cell wall thickness of cambium cells was half as strong compared to cells of the surrounding tissue (Figure 4-figure supplement 1). Assuming that cell wall thickness correlates with cell wall stability, we integrated this feature into our model by expanding VirtualLeaf to allow for the integration of cell-type specific wall stability (see Supporting Information “VirtualLeaf Simulations” for details). We implemented this information in the Hamiltonian operator, which is used to approximate the energy of the system and takes both turgor pressure and cell wall resistance into account. In practice this means that a higher cell wall stability will increase the cell walls’ resistance to being stretched and will result in slower cell* growth.

Utilizing this expanded model (Model 4), we investigated the parameter space to find parameters accurately describing cambium activity not only qualitatively but also quantitatively. To incorporate realistic tissue ratios and unbiased parameter identification, we performed an automated parameter search using a previous characterization of Arabidopsis hypocotyl anatomy [38] as a criterion for parameter selection. To this end, we evaluated our searched parameter sets to aim for a cell type distribution of 24, 10 and 65 % for cambium*, xylem*, and phloem cell* number, respectively. Performing 5000 simulations resulted in n = 5 parameter sets (Supplementary File 2), which produced more realistic cell type proportions than we achieved by our manually selected set before (Figure 4G, Figure 4-figure supplement 2). Thus, by taking real cell type proportions as a guideline for parameter search, we were able to establish a model generating a more realistic morphology as a solution. Furthermore, by generating several parameter sets that described the experimentally observed tissue ratios equally well, we demonstrated that even with differing parameter values the model behavior remains consistent confirming the validity of our approach (Figure 5A, B, Figure 4-figure supplement 2, Video 36, Video 37, Video 38, Video 39, Video 40, Video 41).

**Figure 5.**
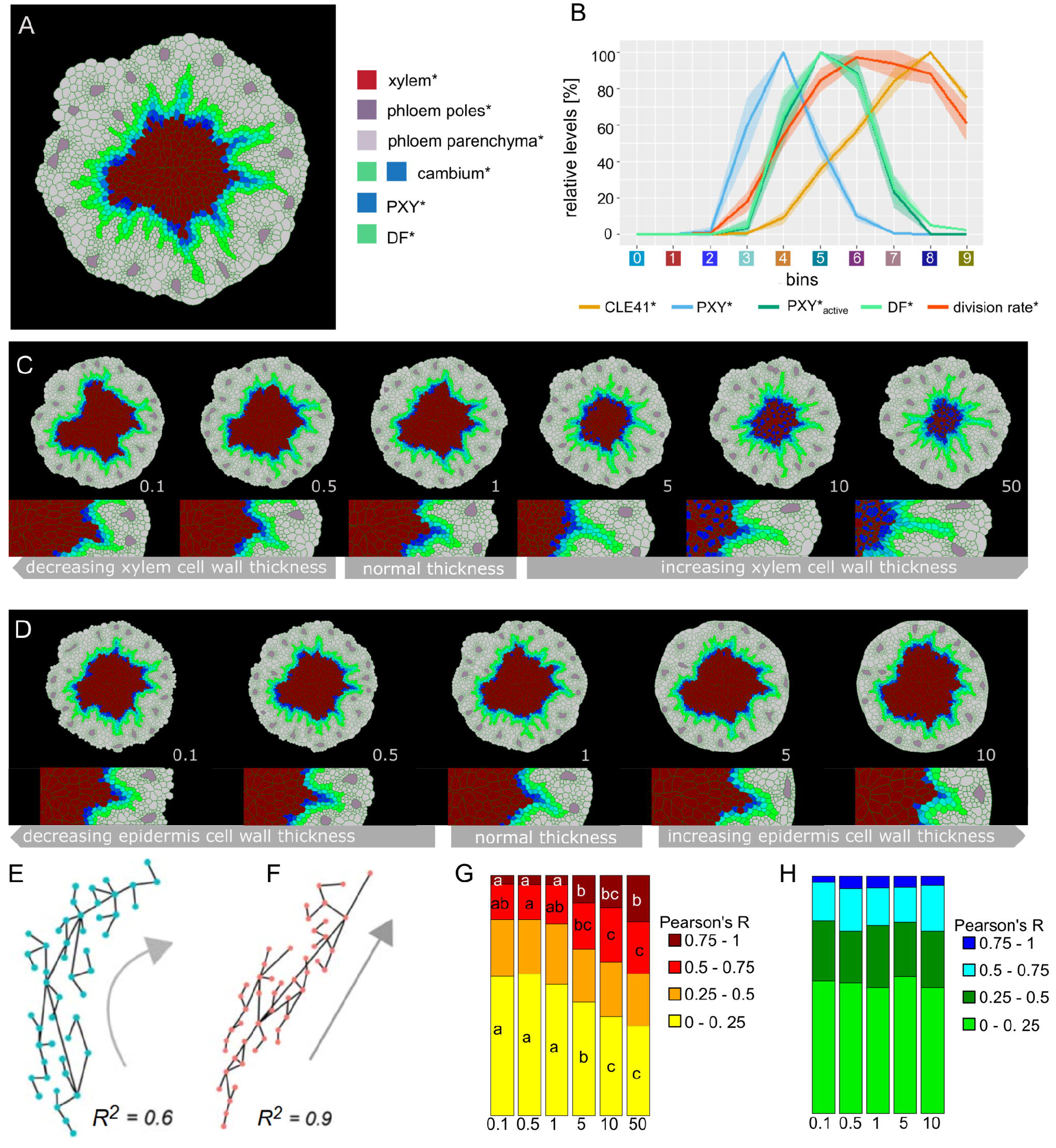
Effect of xylem cell wall thickness* on the radiality of cambium- derived cell lineages*. **(A)** Final output of Model 4 and parameter set 1. **(B)** Visualization of the relative levels of chemicals* and division rates* in different bins. Bin colors along the x-axis correspond to the different bins similarly as in Figure 4C. **(C, D)** Simulation outputs at increasing values of xylem thickness* (C) and epidermis thickness* (D) with the ratio of thickness* vs. experimentally determined xylem thickness indicated at the right bottom corner of each example. All the simulations had the same starting conditions and ran for the same amount of simulated time. At the bottom, there is a magnification of the right region shown in the pictures above, respectively **(E, F)** Examples of the relationship between R^2^ and the geometry of proliferation trajectories (grey arrows) for two different R^2^ values; dots are cell* centroids, lines represent division* events. **(G, H)** Fraction of median relative amount of lineages whose R^2^ falls within a specific range for ten simulations in each condition (n≥70 lineages per simulation) at different xylem thickness* (G) and epidermis thickness* (H) regimes. In case of significant difference among medians, assessed with Kruskal-Wallis (KW significance is *p* < 1.01E-5 for (0, 0.25) interval and *p* < 1.04E-7 for the (0.75,1) interval), the pairwise difference between medians was tested post hoc applying the Dunn test. The post hoc results are reported in each box as letters; medians sharing the same letter or do not display a letter at all do not differ significantly.

To next investigate the role of biomechanics in the direction of cell division, we analyzed the model behavior at different cell wall stability values. Specifically, we were interested in the role of xylem* and epidermis*, the latter being represented by the relative perimeter stiffness of the outer tissue boundary in VirtualLeaf. Of note, defining the outer cell* layer as epidermis* was done for simplicity reasons as the rather complex periderm usually forms the outer tissues of older hypocotyls [47]. Here, we assumed xylem or epidermis cells* and, in turn, the relative perimeter stiffness to be more resistant to expansion due to the thickness of their cell walls* and implemented this behavior in Model 4 as a cellular* property. First we explored how the variation of the thickness of xylem cell walls* impacted tissue formation (Figure 5C, Video 42, Video 43, Video 44, Video 45), we first observed that, as expected, increasing cell wall* thickness led to a xylem*-specific decrease in cell* size and major axis length (Figure 5-figure supplement 1A, B) meaning that xylem cells* not only became smaller but also more isodiametric. In addition, we observed an exclusive decrease in the number of xylem cells* (Figure 5-figure supplement 1D). We explained this effect by a physical constraint generated by ‘stiffer’ xylem cells* acting on neighboring cambium cells* impairing their expansion and, thus, their transformation into xylem* (Video 42, Video 43, Video 44, Video 45, Video 46, Video 47). Importantly, none of these parameters was affected in other cell types* (Figure 5-figure supplement 1), suggesting that the general growth dynamics of the model and especially the behavior of cambium cells* was comparable under the different stiffness* regimes. When analyzing the same characteristics for the different epidermis* tissue regimes (Figure 5D), we found that neither cell size* nor cell length* were impacted (Figure 5-figure supplement 2). Instead, we found a decrease in the number of phloem parenchyma* and phloem pole cells* per simulation with increasing cell wall stability* as the increased resistance of the outer tissue boundary limited the overall tissue growth resulting in less cells being produced in the outer parts of the tissue (Figure 5-figure supplement 2).

To access the effect of increased thickness of xylem and epidermis cell walls* on cell* division orientation, we first defined cell lineages* as groups of cells* having originated from the same precursor cell* and drew lines connecting immediate daughter cells* (Figure 5E, F). We then calculated the goodness of fit (R^2^) of a linear relationship between center of mass coordinates of all individual lines as a proxy for lineage* ‘radiality’ and, thus, for the ratio of periclinal versus anticlinal cell divisions*. After obtaining the R^2^ value for each lineage*, we tested for median differences among r distributions under each thickness regime (Figure 5G, Figure 5-figure supplement 3). These comparisons showed that the increase of the xylem* to non- xylem cell wall* thickness ratio produced a shift from more “curved” lineages (R^2^ < 0.25) towards more radial lineages* (R^2^ > 0.75) (Figure 5G, Figure 5-figure supplement 3A). We attributed this effect to an increased radial elongation of cambium cells* with increasing xylem stiffness* (Video 42, Video 43, Video 44, Video 45, Video 46, Video 47) and the preferred cell division* along the shortest axis in VirtualLeaf [32]. Therefore, implementing thickness as a cell property produced coherent results in terms of the appearance of radial cell files* as an emergent property of cell wall thickness* specifically in the xylem*. In contrast, the analysis of different epidermis cell wall thickness did not show a clear change in the distribution of lineages in the range of analyzed thickness regimes (Figure 5H, Figure 5-figure supplement 3B, Video 48, Video 49, Video 50, Video 51, Video 52) as increasing stiffness regimes limit tissue growth and therefore the formation of cell lineages. These results remained consistent for both xylem* and epidermis* thickness regimes when varying other parameters determining cell wall dynamics, i.e. the target length of cell wall elements and the yielding threshold for the introduction of new cell wall segments (Figure 5-figure supplement 4).

## Discussion

Growth and development of multicellular organisms are complex non-linear processes whose dynamics and network properties are not possible to predict only based on information on their individual building blocks and their one-to-one interactions. The rather simple cellular outline along the radial axes of plant organs, growth in only two dimensions, and the recent identification of central functional properties [11–13], make radial plant growth an attractive target for a systematic approach to reveal its intriguing dynamics. Here, we developed a computational model representing a minimal framework required for radial plant growth using the VirtualLeaf framework [32]. In particular, we combined an agent-based model of the tissue layout with an ODE model of the inter-cellular PXY/CLE41 signaling module. By integrating these two modeling and biological scales, we were able to recapitulate not only the complex behaviors that arise as consequence of the cellular interactions [48] but also the interplay between cellular layout and intercellular signaling dynamics. Therefore, our model allows analyzing fundamental features of plant organ growth and integrates the PXY/CLE41 module as one central element for cambium patterning and maintenance.

Using positional information mediated by morphogenetic gradients of diffusible chemicals to pattern growing structures is a classical concept in developmental biology which has stirred a long history of fundamental debates [49]. Initially, we used the PXY/CLE41 module to generate such a gradient instructing cambium cells* to differentiate into xylem cells*, to proliferate or to differentiate into phloem cells*. Repression of cell division in the distal cambium was achieved by implementing an inhibitory feedback loop of PXY-signaling* on PXY* production. Altogether, this setup was already sufficient to maintain stable radial tissue organization during radial growth and established a maximum of cell division rates in the cambium center as observed by experimental means [13]. Thus, we conclude that cambium organization and radial patterning of plant growth axes can be maintained by a distinct pattern of radially acting morphogens. Such a role was initially proposed for auxin whose differential distribution, however, seems to be rather a result of tissue patterning than being instructive for radial tissue organization [50].

In contrast to expected roles of the *PXY* pathway in xylem formation based on experiments during primary vascular development [19, 22, 24], we observed that the overall amount of proximal tissue production during radial plant growth did not depend on the *PXY* function. Automated image analysis including object classification revealed that neither the number of cells produced toward the organ center nor the number of vessel elements did change in a *pxy* mutant background but rather the ratio between parenchyma and fiber cells. Therefore, in contrast to a negative effect of PXY/CLE41 signaling on vessel formation in vascular bundles in leaves [19, 24], vessel formation during radial plant growth is *PXY/CLE41*- independent. Instead, fiber formation is positively associated with the *PXY/CLE41* module. These observations indicated that xylem formation is unlikely to be instructed by PXY/CLE41 signaling alone and that additional signals are required.

Moreover, the application of markers visualizing cambium organization showed that *PXY*-deficiency leads to cambium disorganization in some regions of the hypocotyl whereas in other areas, cambium anatomy is maintained. Since such areas are regularly spaced, this pattern may arise due to factors acting in parallel to PXY/CLE41 and which also carry spatial information. Although ethylene signaling was reported to act in parallel to PXY/CLE41 signaling, spatial specificity does not seem to be a characteristic property of ethylene signaling [51]. In contrast, PEAR transcription factors are phloem-derived and stimulate the proliferation of cambium stem cells presumably in a PXY/CLE41-independent manner [12] and, thus, may act similarly to the PF* factor we introduced in our model. The ERECTA/EPIDERMAL PATTERNING FACTOR-LIKE (ER/EPFL) receptor-ligand pathway acting in concert with the PXY/CLE41 module [52, 53] represents another candidate for playing such a role. In addition, CLE45 was recently proposed to be expressed in developing sieve elements, the conducting units of the phloem, and repress the establishment of sieve element identity in their immediate environment mediated by the RPK2 receptor protein [39]. The PF* factor in our model combines features of these phloem-derived molecules.

In addition to the phloem sending out instructive signals, early xylem cells have been identified to act as an organizing center of cambium patterning [11]. Although this finding seems to be at odds with our claim that phloem-derived signals are sufficient for cambium organization, it is important to consider that we ignored, for example, upstream regulation of postulated factors like PXY* or CLE41*, which obviously depends on positional information. For simplicity, we also ignored organizing effects of signaling longitudinally to cross sections as it can, for example expected for polar auxin transport [54–56] in the context of cambium activity or xylem formation. Although being considerably more complex, the establishment of 3D models will be crucial essential for addressing this aspect.

In this context, it is interesting to note that we deliberately excluded the transition from the initially bisymmetric tissue conformation to a concentric tissue organization as it occurs in hypocotyls and roots [11, 38] from our model. Our rationale was that the rather complex change in tissue anatomy from a primary to a secondary conformation in the hypocotyl required more assumptions in our model and would have spoiled the advantages of a relatively simple anatomy for generating a cell-based computational model. Moreover, the differences in primary anatomy of shoots and roots before the onset of radial plant growth [11, 57] would have required different cellular outlines for both cases and, thus, would have hampered the generality of our approach.

Interestingly, the front of cambium domains is very stable, i.e. almost perfectly circular, *in planta* but this is not the case for our computational simulations. We believe that instability in the computational models is due to local noise in the cellular pattern leading to differential diffusion of chemicals* with respect to their radial position and to a progressive deviation of domains from a perfect circle. Such a deviation seems to be corrected by an unknown mechanism *in planta* but such a corrective mechanism is, due to the absence of a good indication *in planta*, not implemented in our models. Time course analyses of anatomies of wt and *pxy* lines (Figure S2) revealed ‘gaps’ in the cambium domain already at early stages of *pxy* development arguing against the possibility that the *pxy* anatomy is caused by increased front instability. Although a corrective mechanism ensuring front stability *in planta* is difficult to predict, our model now allows to test respective ideas like directional movement of chemicals or stabilizing communication between cells during cambium activity. In this context it is interesting that increasing epidermis* ‘stiffness’ increased circularity of the growing organ* which may be administered by the periderm [47], the protective cell layers which we did not consider in our model.

Current research on plant mechanical biology indicates how cell mechanical properties influence cell and tissue morphogenesis. Microtubules, turgor pressure and cell wall composition are central factors in this regard [58, 59]. Due to the geometric constraints in a radially growing plant axis, it becomes challenging to uncouple these factors experimentally and to establish the impact of one factor on organ patterning during radial plant growth. By expanding VirtualLeaf to allow for the integration of cell type-specific wall stability, we fundamentally increased the spectrum of potential modeling approaches. In particular, since cell wall stability is accessible by the cellular model throughout simulations, it is now possible to simulate and analyze e.g. the dynamics of auxin or brassinosteroid-mediated cell wall loosening [60, 61]. In our cambium model, by modulating exclusively cellular ‘stiffness’, we were able to computationally simplify the ‘physical’ properties and, thereby, develop a hypothesis how inter-tissue forces influence stem cell behavior not only cell autonomously, but also in a non-cell autonomous manner.

Taken together, we envision that the model presented in this study recapitulates the qualitative and quantitative variation in radial plant growth on multiple levels, found in different mutants or when comparing different dicotyledonous species [62]. Remarkable features like the establishment of concentric cambium rings often found in the order of *Caryophyllales* [63] or ‘phloem wedges’ found in the *Bignonieae* genus [64] may be recapitulated by adjusting the model’s parameters values or by introducing additional factors. In the future, the model may help to predict targets of environmental stimuli inducing changes of cambium activity like seasonal changes [64] or mechanical perturbation [66], allowing the generation of testable hypotheses.

Thus, our dynamic model will be a useful tool for investigating a process not possible to observe in real time and partly develops over exceptionally long periods.

## Material and Methods

### Plant material and growth conditions

*Arabidopsis thaliana* (L.) Heynh. plants of Columbia-0 accession were used for all experiments and grown as described previously [23]. *pxy-4* (SALK_009542, N800038) mutants were ordered from the Nottingham Arabidopsis Stock Centre (NASC). Plant lines carrying *IRX3pro:CLE41* and *35Spro:CLE41* transgenes [16] were kindly provided by Peter Etchells (Durham University, UK). *PXYpro:ECFP-ER* (*pPS19*) and *SMXL5pro:EYFP-ER* (*pJA24*) reporter lines expressing fluorescent proteins targeted to the endoplasmatic reticulum (ER) were described previously [67, 68].

### Confocal microscopy

Hypocotyls were isolated and cleaned from surrounding leaf material using razor blades (Classic Wilkinson, Germany). The cleaned hypocotyls were mounted in 7 % low melting point agar (Sigma-Aldrich, St. Louis, MO, USA) in water and sections were generated using a vibratome (Leica VT1000 S). For Figure 4-figure supplement 1, 75 µm thick sections were stained for 60 minutes with 0.1 % w/v Direct Yellow 96 (Sigma Aldrich, St. Louis, US, S472409-1G) diluted in ClearSee [46] (10 % w/v xylitol, 15 % w/v sodium deoxycholate, 25 % w/v urea), washed three times with ClearSee and mounted in ClearSee on microscope slides. For other experiments, 60 μm thick sections were stained for 5 minutes with 0.1 % w/v Direct Red 23 (Sigma Aldrich, St. Louis, US, 212490-50G) diluted in water, washed three times with water and mounted in water on microscope slides. For analyzing the fluorescent markers, a Leica SP8 or Stellaris 8 (Leica, Germany) confocal microscope was used. Different fluorescence protein signals were collected in different tracks. YFP was excited at 514 nm and emission was collected at 522-542 nm. CFP was excited at 458 nm and the signal emission was collected at 469-490 nm. The Direct Red 23-derived signal was excited at 495 nm and emission was detected at 558-649 nm. The Direct Yellow 96-derived signal was excited at 488 nm and emission was detected at 500-540 nm. For qualitative comparisons, 5-10 samples for each sample type were included and repeated at least twice.

### Ilastik cell type counting

For cell type classification and quantification, sections were produced as previously described [23]. The xylem area was cropped manually from histological images of wild type and *pxy* mutant. The Ilastik toolkit [69] was used for image segmentation and cell type classification (https://www.ilastik.org). With a training set, the pixel classification workflow was trained to distinguish cell walls from the background. After segmentation, the object classifier was then trained to split the resulting objects into four groups - xylem vessels, xylem fibers, xylem parenchyma, and unclassified objects. The resulting classifier was then applied to all cropped images. For each image, cell type data were extracted using python. 11-12 plants each for wild type and *pxy* mutants were compared in two independent experiments.

### VirtualLeaf simulations

Simulations were performed as recommended previously [32]. To be able to see established models in action, the VirtualLeaf software was installed according to the following instructions described in the Appendix 1 and as described previously [70]. All simulations within Model 1, Model 2, Model 3, and Model 4 respectively, were conducted for the same VirtualLeaf time duration and repeated at least ten times to account for the stochastic nature of the tissue simulations (for details on simulations in VirtualLeaf see section “Description of the VirtualLeaf simulations” in the Appendix 1. Information on Supplementary Items and Videos are found in the Supplementary Information Items file.

### Splitting the result of VirtualLeaf simulations into bins

After a VirtualLeaf simulation was completed, the resulting xml template was stored. To analyze the distribution of chemicals* in such a template along the radial axis, we produced a python script named “Cambium_bins_calculation.ipnb”. Within the script, it was possible to indicate the path to the xml file, and the script produced two .csv files – one with a table containing data about each cell and another with information about averages across the requested bin number.

### Parameter estimation and exploration of the parameter space

To estimate the model parameters and, at the same time, investigate the parameter space, we performed a large set of simulations with randomized parameters to identify feasible parameter combinations. In particular, we employed a combination of python and shell scripting to set up the parameter sets, run the simulations and analyze the results. To generate the parameter sets we followed the tutorial using the python library xml.etree.Elementree as described [33]. The search intervals were defined based on the manually determined parameter values of model 3A: The search interval was set between 1/3 and 3-times the original value. The individual parameter sets were then simulated for a duration of t_simulated = 2200 steps on a computing cluster (Linux, 64-bit). The resulting xml leaf was then analyzed based on tissue size and proportions. Based on *in planta* observations [38], we determined that the simulation should result in 24 % cambium, 10 % xylem, and 65 % phloem cells.

As all tissues are equally important, we used a weighted least squares scoring function to compare the experimentally measured tissue ratios with the model simulations. We added a term for the total number of cells to favor parameter sets that resulted in tissue growth. Altogether, this resulted in the following scoring function:

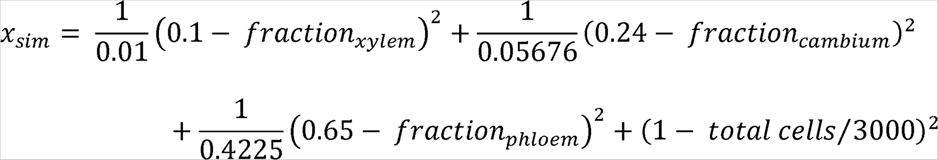

As we were interested in obtaining simulations with an active cambium we discarded simulations that resulted in hypocotyls* with less than 300 cells* in total and with cambium cells less than 30.

### Exploration of stiffness

To explore the effects of stiffer (i.e. less flexible) xylem cell walls and epidermis cell walls as represented by the perimeter stiffness, we slightly modified the VirtualLeaf code so that it was possible for λ_L_ (the “cost” of deviation of the wall element’s length from the target length) to assume cell type-specific values. More specifically, we defined a new parameter named cell wall stablity, and set λ_L_ = cell wall stablity according to the experimentally determined cell wall thickness as a proxy for cell wall stability. We then ran the model with different ratios of cell wall stablity compared to the normal parameter value, while maintaining the same tissue configuration used for the other simulations done within this study. The values chosen for the parameter were 0.1, 0.5, 1, 5, 10-fold change for both tissues of interest and 50-fold change for xylem*. We replicated each run 30 times. We further repeated the analysis of different thickness regimes while varying other cell wall dynamic parameters by +/- 50%, i.e. the target element for cell wall elements and the yielding threshold for the introduction of new cell wall elements (for n=10 simulations per parameter combination).

To study the proliferation trajectory of cells, we performed for every lineage a linear regression of the centers of mass for the cells belonging to that lineage, and used the coefficient of determination (R^2^) as proxy for proliferation trajectory of the lineage. We next tested for median differences among the R^2^ distribution under each thickness regime using the Kruskal-Wallis (KS) test, and performed the Dunn test to determine differences among groups in case of significant KS. Before performing the KS, we subsampled the data to maintain the same number of samples across thickness values, and bootstrapped the samples to obtain robust median estimators and confidence intervals.

Statistical analyses and visualizations of ‘stiffness’ were performed using the R language for statistical computing and graphics (https://www.r-project.org/), using the tidyverse family of packages [71], together with the broom (https://cran.r-project.org/web/packages/broom/index.html), FSA (https://github.com/droglenc/FSA), and boot packages [72, 73].

## Material availability statement

For each of the models and frameworks described in this paper, we provide code files at https://github.com/thomasgreb/Lebovka-et-al_cambium-models.

## Supporting information

Supplementary Information Items

## Acknowledgements

We thank Peter Etchells (Durham University, UK) for providing seed material, Karin Grünwald and Martina Laaber-Schwarz (both GMI, Vienna, Austria) for technical assistance and Dongbo Shi, Eva-Sophie Wallner and Vadir López-Salmerón for comments on the experimental strategy and the manuscript. We also thank Claudiu Antonovici (University of Leiden, The Netherlands) for help in setting up the VirtualLeaf platform. This work was supported by the Deutsche Forschungsgemeinschaft (DFG) through the Research Unit FOR2581 ‘Plant Morphodynamics, grant GR2104/4-1 and a Heisenberg Professorship (GR2104/5-2) to T.G.. The work by B.H.M. was initiated at Centrum Wiskunde & Informatica (CWI), Amsterdam, The Netherlands. R.M. and B.H.M. thank CWI for providing a CWI Internship to B.H.M. and for hosting I.L.. R.G. was supported by the CRC 1101 „Molecular Encoding of Specificity in Plant Processes“ (DFG) and the Joachim Herz Stiftung.

## Conflict of Interest

The authors have no conflicts of interest to declare.

## Supplementary Information Items

**Figure 3-figure supplement 1:** Magnification of results of cell type classification shown in Figure 3.

**Figure 3-figure supplement 2:** Dynamics of *PXYpro:CFP/SMXL5pro:YFP* activities during radial hypocotyl growth in wild type, *IRX3pro:CLE41* and *pxy* plants.

**Figure 4-figure supplement 1:** Determination of cell wall thickness across the radial sequence of hypocotyl tissues.

**Figure 4-figure supplement 2:** Behavior of the different model parameterizations (Model 4:2-5)

**Figure 5-figure supplement 1:** Distribution of cell* properties under different xylem ‘stiffness’ regimes.

**Figure 5-figure supplement 2:** Distribution of cell* properties under different tissue boundary (=epidermis*) ‘stiffness’ regimes.

**Figure 5-figure supplement 3:** Fraction of median relative amount of cell lineages for parameter sets 2-5.

**Figure 5-figure supplement 4:** Fraction of median relative amount of cell lineages at different parameters governing cell wall* dynamics.

**Supplementary File 1:**Table listing cell* behavior rules for Models 1-4.

**Supplementary File 2:**Table listing parameter values and chemicals.

**Appendix 1:** Instructions for implementing VirtualLeaf models.

**Video 1:** Model 1 output, visualizing xylem (red) and phloem (purple), and accumulation of PXY* (blue) and PXY_active_* (green)

**Video 2:** Model 1 output, visualizing CLE41* (yellow) accumulation

**Video 3:** Model 1 output, visualizing cell divisions (red)

**Video 4:** Model 1 output, visualizing PXY_active_*

**Video 5:** Model 1 output, visualizing PXY*

**Video 6:** Model 2A output, visualizing xylem (red) and phloem (purple), and accumulation of PXY* (blue) and PXY-active* (green)

**Video 7:** Model 2A output, visualizing CLE41* (yellow) accumulation

**Video 8:** Model 2A output, visualizing cell divisions (red)

**Video 9:** Model 2A output, visualizing cell divisions (red) together with PXY* (blue) and PXY-active* (green) accumulation

**Video 10:** Model 2A output, visualizing PXY_active_*

**Video 11:** Model 2A output, visualizing PXY*

**Video 12:** Model 2B output, visualizing xylem (red) and phloem (purple), and accumulation of PXY* (blue), and PXY-active* (green)

**Video 13:** Model 2B output, visualizing CLE41* (yellow) accumulation

**Video 14:** Model 2B output, visualizing cell divisions (red)

**Video 15:** Model 2B output, visualizing accumulation of PXY* (blue) and PXY- active* (green)

**Video 16:** Model 2B output, visualizing PXY_active_*

**Video 17:** Model 2B output, visualizing PXY*

**Video 18:** Model 3A output, visualizing xylem (red), phloem parenchyma (light purple), and phloem poles (dark purple), and accumulation of PXY* (blue) and the division chemical (DF)* (green)

**Video 19:** Model 3A output, visualizing CLE41* (yellow) accumulation

**Video 20:** Model 3A output, visualizing cell divisions (red)

**Video 21:** Model 3A output, visualizing accumulation of PXY* (blue) and PXY_active_* (green)

**Video 22:** Model 3A output, visualizing PXY_active_*

**Video 23:** Model 3A output, visualizing PXY*

**Video 24:** Model 3B output, visualizing xylem (red), phloem parenchyma (light purple), and phloem poles (dark purple), and accumulation of PXY* (blue) and the division chemical (DF)* (green)

**Video 25:** Model 3B output, visualizing CLE41* (yellow) accumulation

**Video 26:** Model 3B output, visualizing cell divisions (red)

**Video 27:** Model 3B output, visualizing accumulation of PXY* (blue) and PXY_active_* (green)

**Video 28:** Model 3B output, visualizing PXY_active_*

**Video 29:** Model 3B output, visualizing PXY*

**Video 30:** Model 3C output, visualizing xylem (red), phloem parenchyma (light purple), and phloem poles (dark purple), and accumulation of PXY* (blue) and the division chemical (DF)* (green)

**Video 31:** Model 3C output, visualizing CLE41* (yellow) accumulation

**Video 32:** Model 3C output, visualizing cell divisions (red)

**Video 33:** Model 3C output, visualizing accumulation of PXY* (blue) and the division chemical (DF)* (green)

**Video 34:** Model 3C output, visualizing PXY_active_*

**Video 35:** Model 3C output, visualizing PXY*

**Video 36:** Model 4 output, parameter Set 1, visualizing xylem (red), phloem parenchyma (light purple), and phloem poles (dark purple), and accumulation of PXY* (blue) and the division chemical (DF)* (green)

**Video 37:** Model 4 output, parameter set 1, visualizing CLE41* (yellow) accumulation

**Video 38:** Model 4 output, parameter set 1, cell divisions (red)

**Video 39:** Model 4 output, parameter set 1, visualizing accumulation of PXY* (blue) and the division chemical (DF)* (green)

**Video 40:** Model 4 output, parameter set 1, visualizing PXY_active_*

**Video 41:** Model 4 output, parameter set 1, visualizing PXY*

**Video 42:** Model 4 output, visualizing accumulation of PXY* (blue) and the division chemical (DF)* (green) implementing a 0.1-fold change in xylem* cell wall stiffness

**Video 43:** Model 4 output, visualizing accumulation of PXY* (blue) and the division chemical (DF)* (green) implementing a 0.5-fold change in xylem* cell wall stiffness

**Video 44:** Model 4 output, visualizing accumulation of PXY* (blue) and the division chemical (DF)* (green) at experimentally determined xylem cell wall thickness/stability

**Video 45:** Model 4 output, visualizing accumulation of PXY* (blue) and the division chemical (DF)* (green), implementing a 5-fold increase in xylem* cell wall stiffness

**Video 46:** Model 4 output, visualizing accumulation of PXY* (blue) and the division chemical (DF)* (green), a 10-fold increase in xylem* cell wall stiffness

**Video 47:** Model 4 output, visualizing accumulation of PXY* (blue) and the division chemical (DF)* (green), a 50-fold increase in xylem* cell wall stiffness

**Video 48:** Model 4 output, visualizing accumulation of PXY* (blue) and the division chemical (DF)* (green) implementing a 0.1-fold change in epidermis* cell wall stiffness

**Video 49:** Model 4 output, visualizing accumulation of PXY* (blue) and the division chemical (DF)* (green) implementing a 0.5-fold change in epidermis* cell wall stiffness

**Video 50:** Model 4 output, visualizing accumulation of PXY* (blue) and the division chemical (DF)* (green) at experimentally determined epidermis* cell wall thickness/stability

**Video 51:** Model 4 output, visualizing accumulation of PXY* (blue) and the division chemical (DF)* (green), implementing a 5-fold increase in epidermis* cell wall stiffness

**Video 52:** Model 4 output, visualizing accumulation of PXY* (blue) and the division chemical (DF)* (green), a 10-fold increase in epidermis* cell wall stiffness

## Notes

https://github.com/thomasgreb/Lebovka-et-al_cambium-models

### Competing Interest Statement

The authors have declared no competing interest.

### Summary of Updates

In the revised manuscript, we provide a new VirtualLeaf version in which the modulation of cell wall mechanics is a general optional feature: Cell specific cell wall stability values can be accessed throughout simulations and allow for a more precise description of the mechanics effects of cell differentiation and growth, which opens up new opportunities for computational analyses. Moreover, we provide expanded mathematical descriptions, parameter analysis of our models, and a time course analysis of related hypocotyl anatomies. Supplementary material has been expanded and now contains a determination of cell wall thickness and extended model characterizations. In comparison to the first version, we added the following authors: Xiaomin Liu generated data shown in Figure 3-figure supplement 2, Theresa Schlamp generated data shown in Figure 4-figure supplement 1, and Ruth Grosseholz advanced computational models, redid all model simulations and generated respective figures and videos.

