## Supplementary material for "Computational modelling of cambium activity provides a regulatory framework for simulating radial plant growth": Figure 3-figure supplement 1

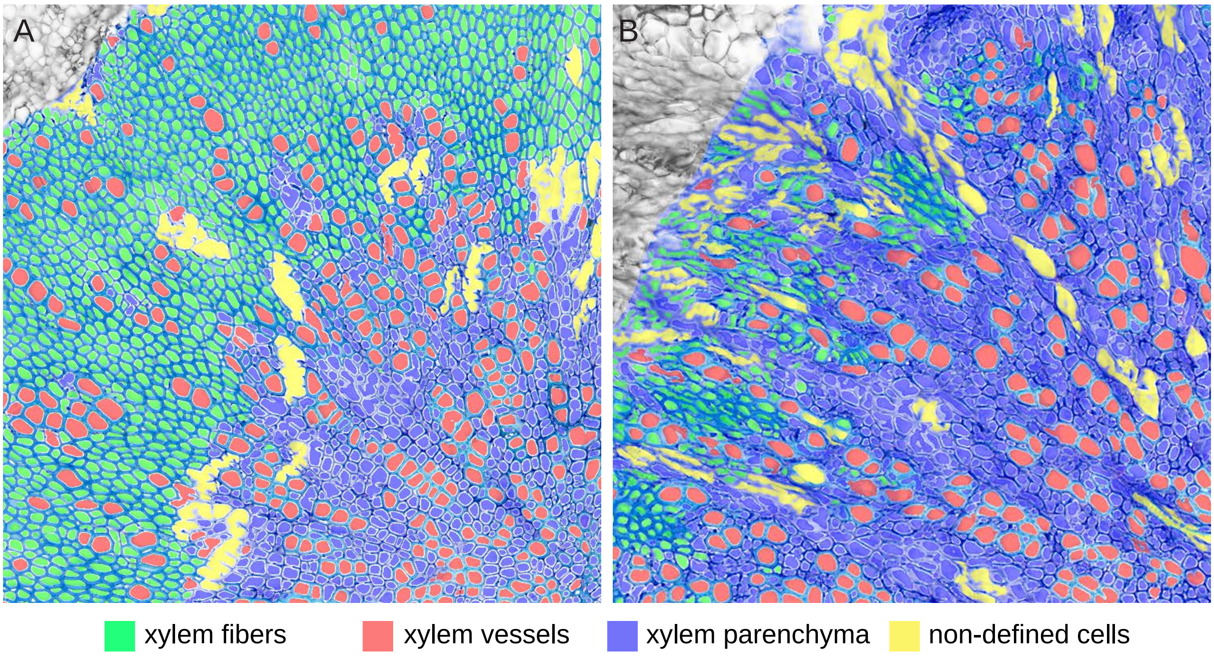


**Figure 3—figure supplement 1: Magnification of results of cell type classification shown in Fig. 3.**
(A) wild type
(B) *pxy* mutant
