## Supplementary material for "Computational modelling of cambium activity provides a regulatory framework for simulating radial plant growth": Figure 3-figure supplement 2

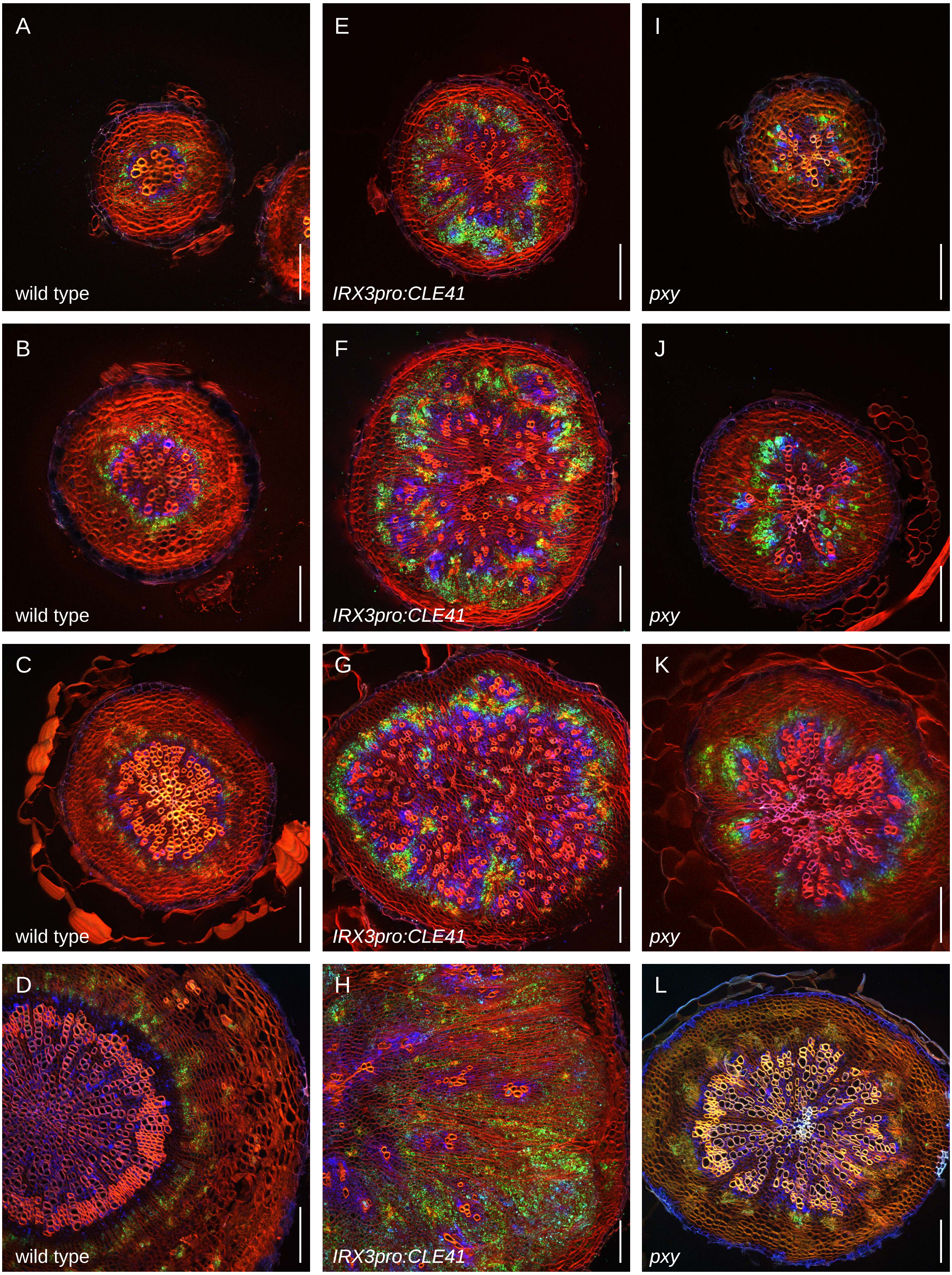


**Figure 3—figure supplement 2. Dynamics of *PXYpro:CFP/SMXL5pro:YFP* activities during radial hypocotyl growth in wild type, *IRX3pro:CLE41* and *pxy* plants.**
**(A-D)** *PXYpro:CFP* (blue) and *SMXL5pro:YFP* (green) activities at different stages of wild type hypocotyl development from young (A) to old (D). **(E-H)** *PXYpro:CFP* (blue) and *SMXL5pro:YFP* (green) activities at different stages of hypocotyl development in *IRX3pro:CLE41* plants from young (A) to old (D). **(I-L)** *PXYpro:CFP* (blue) and *SMXL5pro:YFP* (green) activities at different stages of hypocotyl development in *pxy* mutants from young (A) to old (D). Sections are stained by Direct Red 23 (red). Scale bars: 100 µm. Note that pictures D, H and L are also depicted in Figure 2 and Figure 3.
