## Supplementary material for "Computational modelling of cambium activity provides a regulatory framework for simulating radial plant growth": Figure 4-figure supplement 1

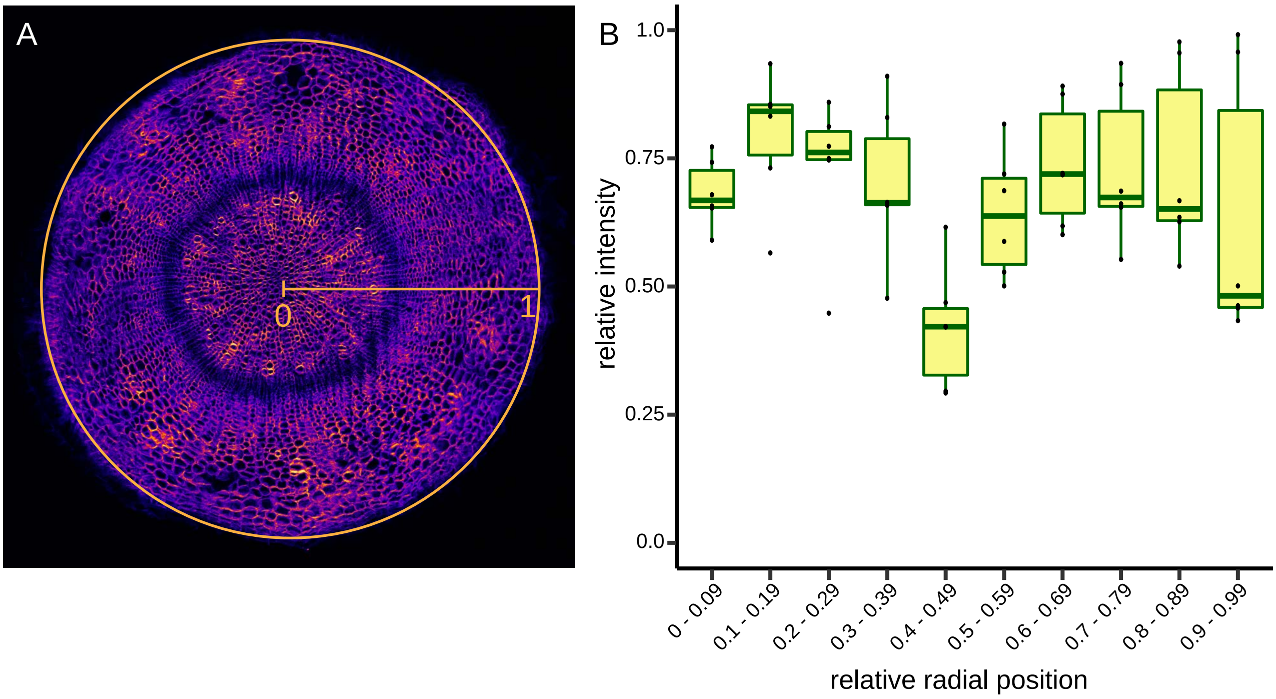


**Figure 4—figure supplement 1: Determination of cell wall thickness across the radial sequence of hypocotyl tissues.**

**(A)** Cross section of a 4.5 week-old plant stained by Direct Yellow 96. Radius and circumference used by the 'Radial Profile' function of the Fiji image analysis tool [72] is indicated. Note that the function uses the whole circle area for analysis.

**(B)** Plot of staining intensities from six Direct Yellow 96-stained cross sections analyzed by the 'Radial Profile' function of Fiji. Staining intensity and radius were normalized to 1 by dividing obtained values by maximum values within respective sample data sets. Intensity profiles were binned in 10 equal parts and the median intensity of single samples was calculated, indicated as dots. The boxplot shows the variation among the samples and the mean of the dataset of each bin.
