## Supplementary material for "Computational modelling of cambium activity provides a regulatory framework for simulating radial plant growth": Figure 4-figure supplement 2

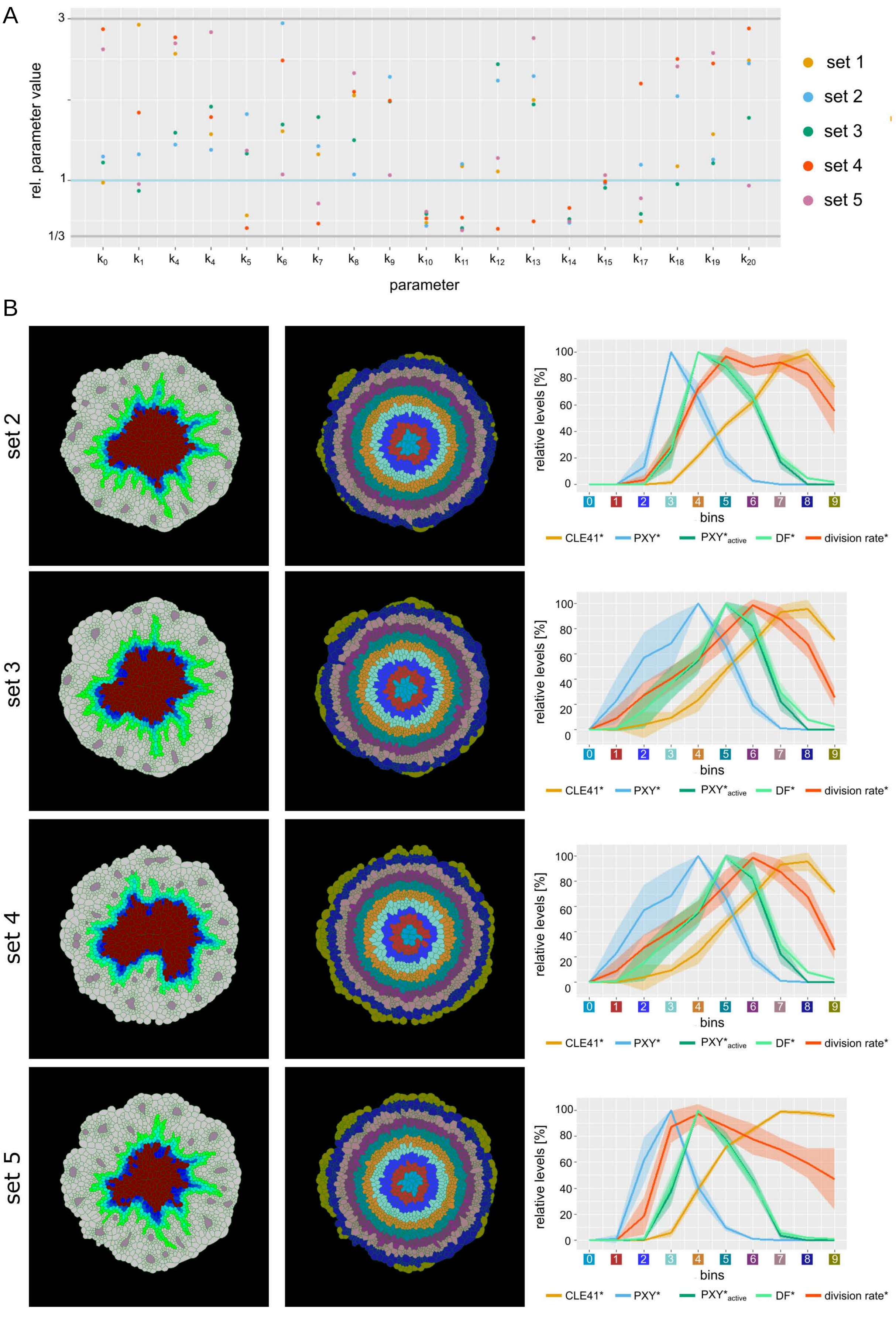


**Figure 4—figure supplement 2: Behavior of the different model parameterizations (Model 3D:2-5).**

**(A)** Overview of parameter values of the different parameter sets. Shown are the relative values of the estimated parameter compared to the original parameter values. Horizontal lines indicate the lower (1/3) and upper (3-fold) boundary (grey) as well as the original parameter value (blue).

**(B)** Behavior of parameter sets 2-5. Shown is the final output of the simulation, the tissue* sorted into bins as well as the average chemical concentration* per bin (for n=10 simulations).
