## Supplementary material for "Computational modelling of cambium activity provides a regulatory framework for simulating radial plant growth": Figure 5-figure supplement 1

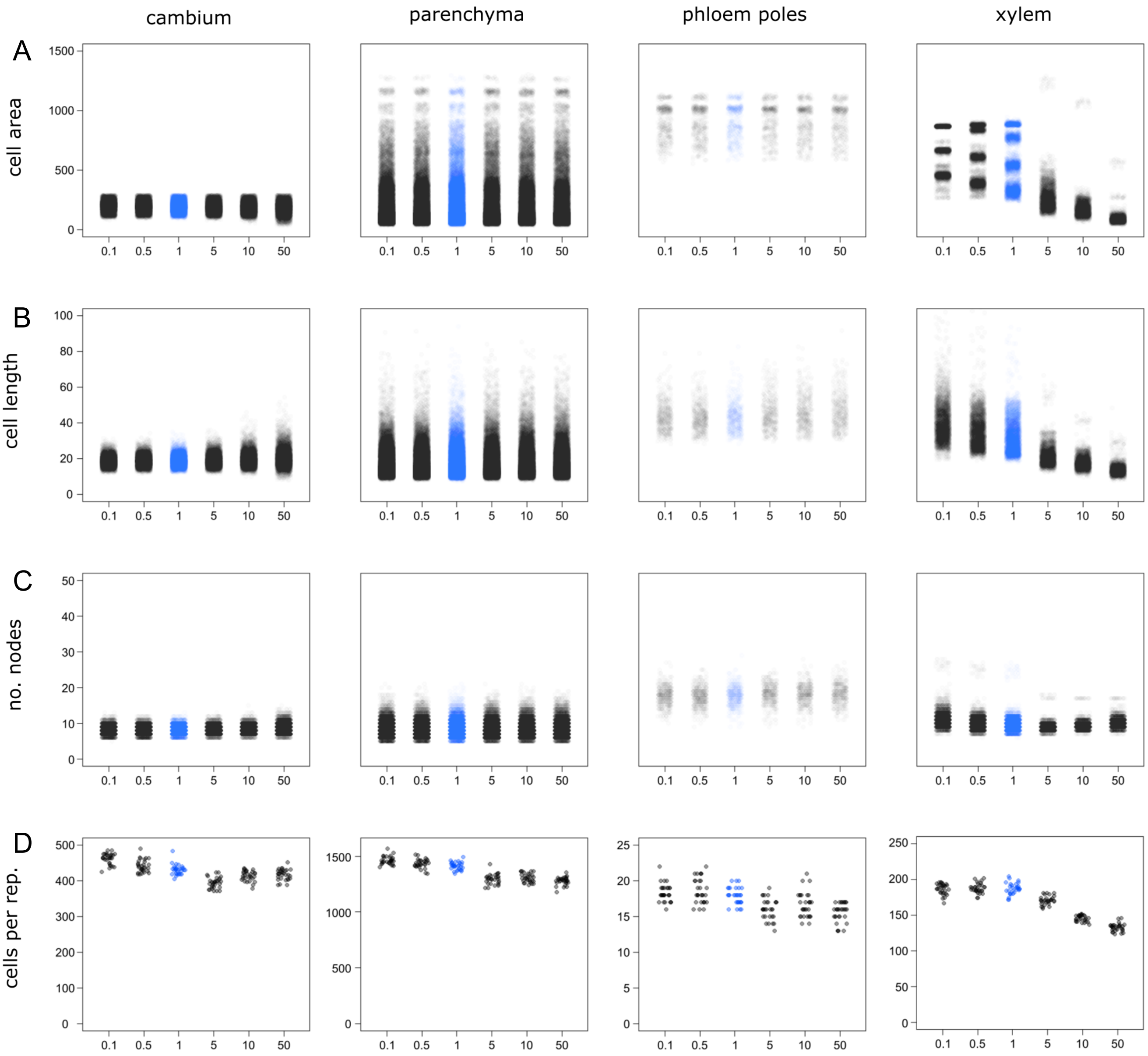


**Figure 5—figure supplement 1: Distribution of cell* properties under different xylem ‘stiffness’ regimes**
**A)** Cell* size in arbitrary units
**B)** Major axis lengths of cells* in arbitrary units
**C)** Numbers of nodes (vertexes) per cell*
**D)** Numbers of cells*
among cell types and thickness values for n = 30 simulations under each thickness regime. The blue color highlights the simulation at the experimentally determined thickness value. The x-axis indicates values of xylem thickness as the ratio of xylem thickness* vs. experimentally determined thickness. A slight horizontal displacement of points has been added to enhance visualization. Values for individual cells* found in all 30 simulations are displayed in A-C, whereas numbers of cells* in each cell type* for each one of the 30 simulations are shown in D.
