## Supplementary material for "Computational modelling of cambium activity provides a regulatory framework for simulating radial plant growth": Figure 5-figure supplement 2

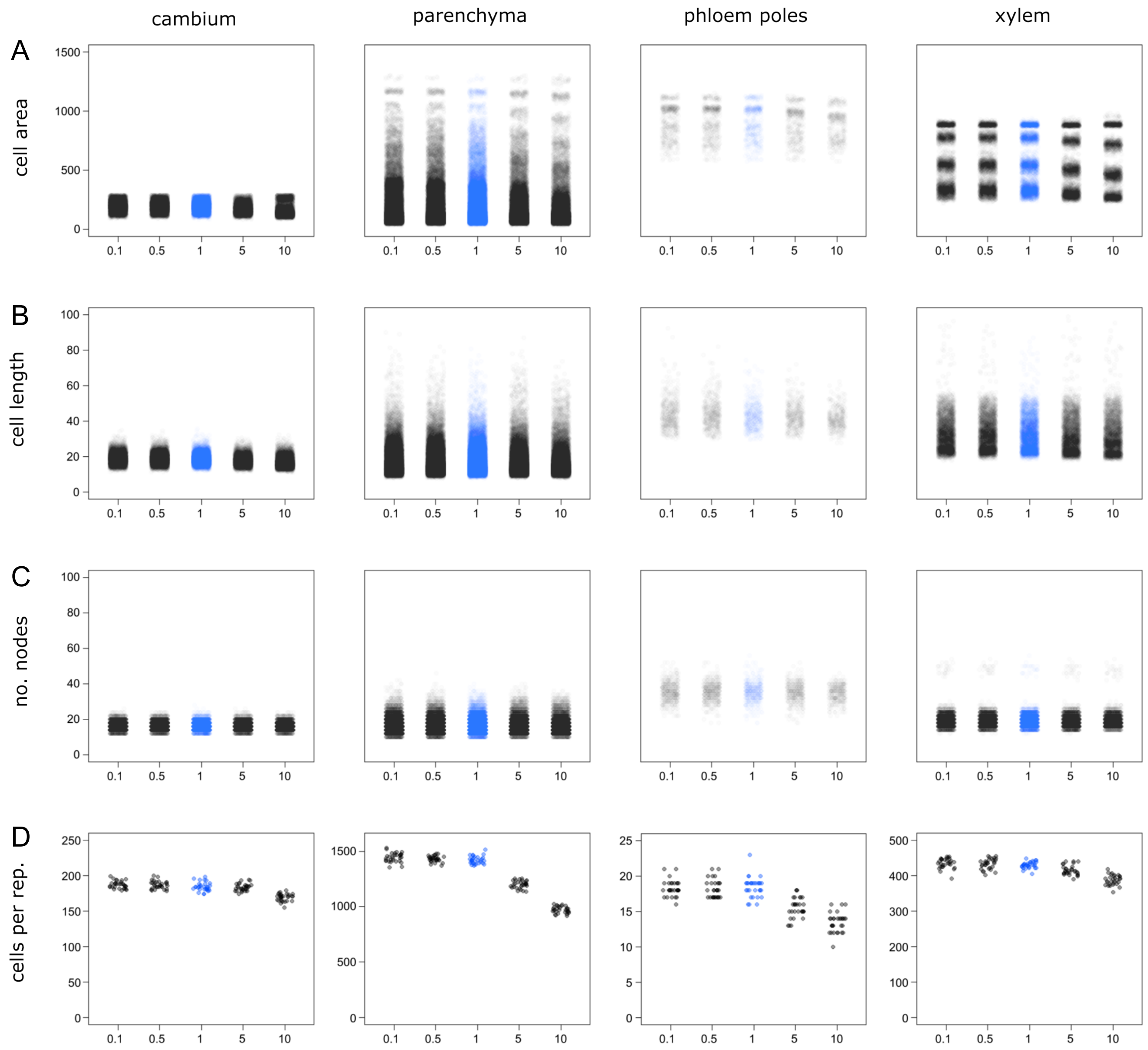


**Figure 5—figure supplement 2: Distribution of cell* properties under different tissue boundary (=epidermis*) ‘stiffness’ regimes**

**(A)** Cell* size in arbitrary units

**(B)** Major axis lengths of cells* in arbitrary units

**(C)** Numbers of nodes (vertexes) per cell*

**(D)** Number of cells*

among cell types and thickness values for n = 30 simulations under each thickness regime. The blue color highlights the simulation running at normal thickness level; the x-axis indicates values of the relative perimeter thickness as the fold-change compared to the standard parameters. A slight horizontal displacement of points has been added to enhance visualization. Values for individual cells* found in all 30 simulations are displayed in A-C, whereas numbers of cells* in each cell type* for each one of the 30 simulations are shown in D.
