## Supplementary material for "Computational modelling of cambium activity provides a regulatory framework for simulating radial plant growth": Figure 5-figure supplement 3

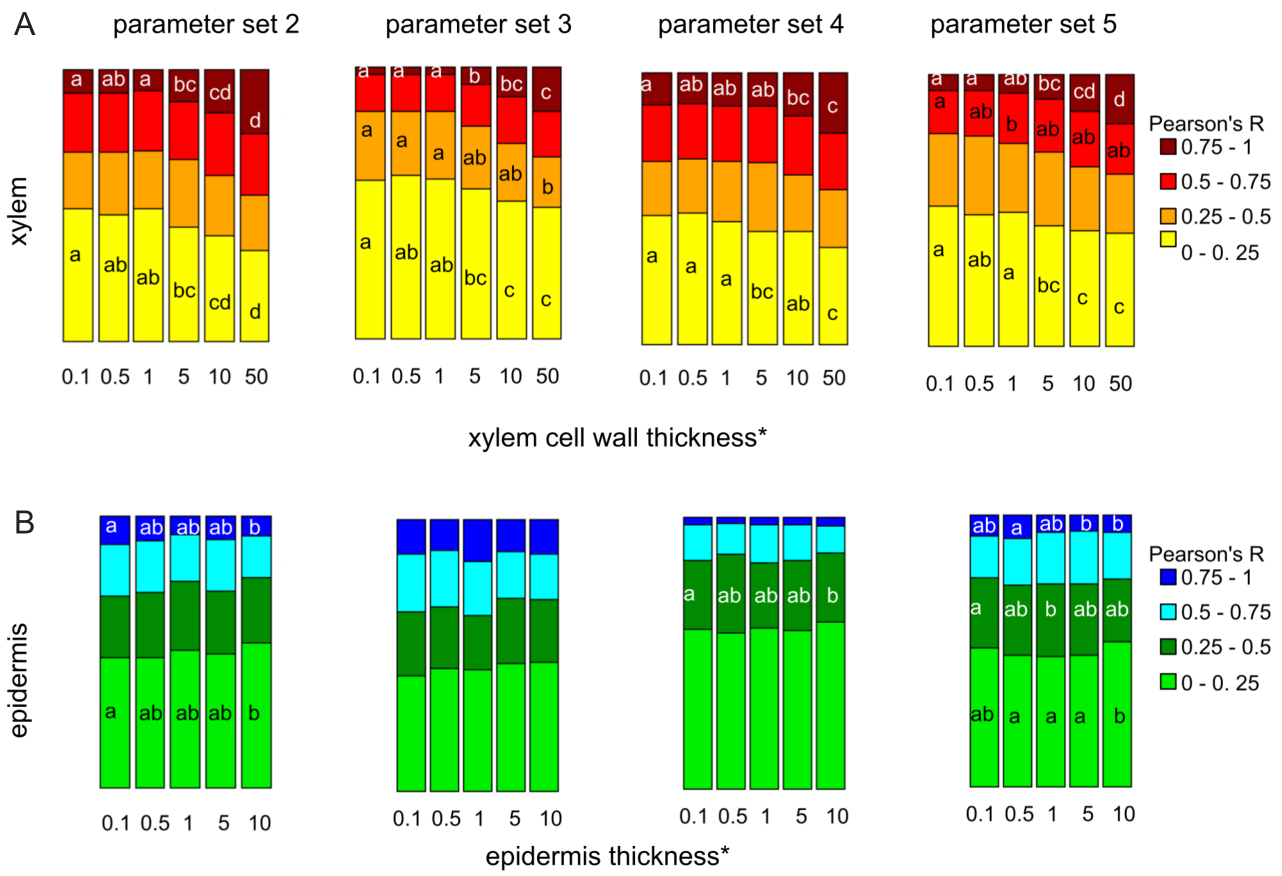


**Figure 5—figure supplement 3: Fraction of median relative amount of cell lineages for parameter sets 2-5.**
**(A)** With increasing xylem* ‘thickness’
**(B)** With increasing epidermis* ‘thickness’
