## Supplementary material for "Computational modelling of cambium activity provides a regulatory framework for simulating radial plant growth": Figure 5-figure supplement 4

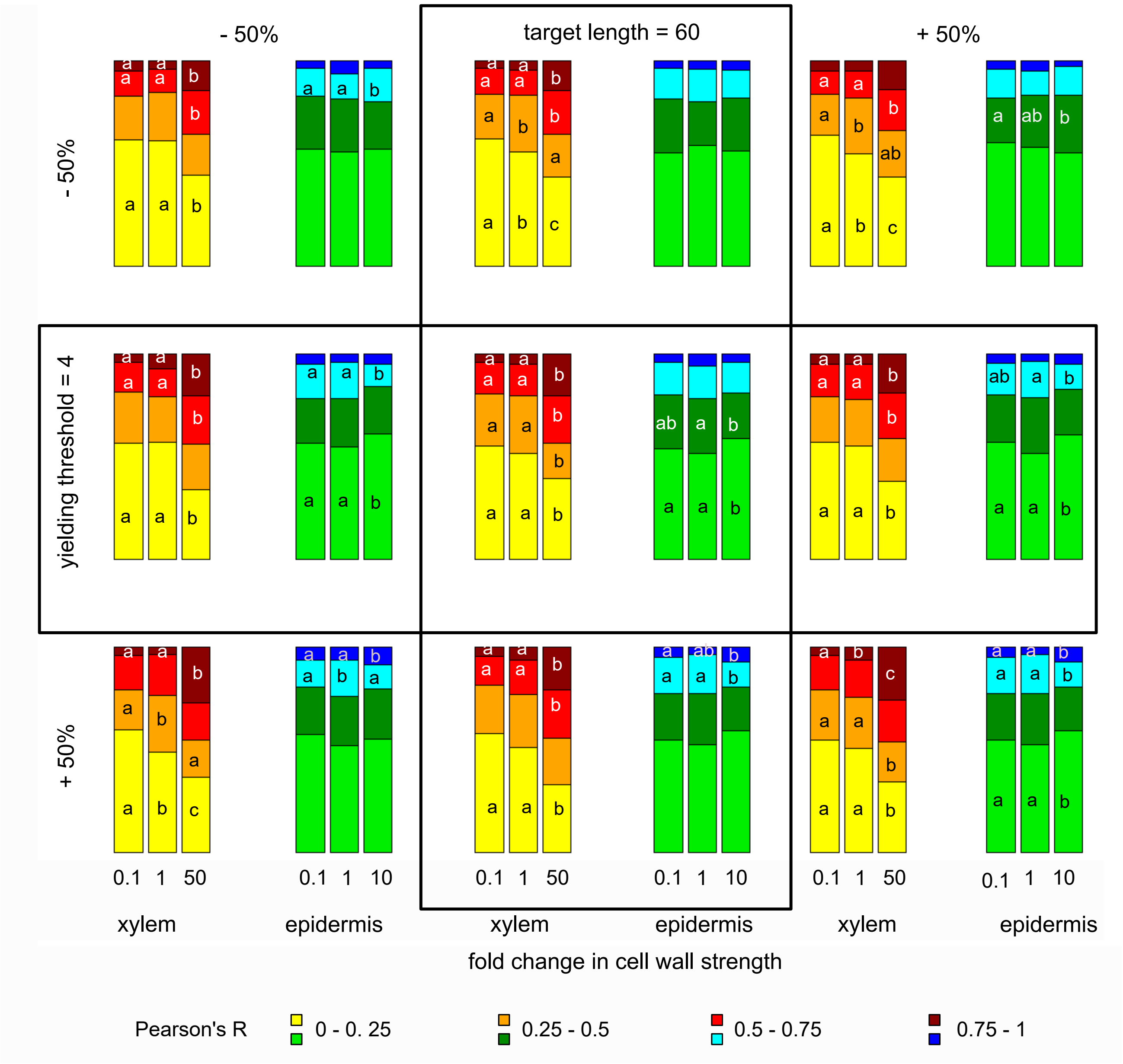


**Figure 5—figure supplement 4: Fraction of median relative amount of cell lineages at different parameters governing cell wall* dynamics.** The model parameters cell walls' target length and yielding threshold were varied by +/-50% and the behavior at different cell wall stability values simulated. The statistical analysis was done as described before for Fig. 5 for n=10 simulations each and n≥70 cell lineages per simulation.
