## Supplementary File 1 for "Computational modelling of cambium activity provides a regulatory framework for simulating radial plant growth"

**Overview of the cell behavior rules for the different cell types in the different model structures.** Unless otherwise specified all cell behaviors take place as long as all conditions listed for the behavior remain true. For special cases (the differentiation to phloem in Model 2 as well as the differentiation to phloem and xylem in Models 3 &4) these only take place instead of the previous cell behavior rules. The cell behavior rules listed for one model structure apply to all variations of that model, e.g. the cell behaviors and condition for Model 2 are identical for Model 2 A, B, C and D. The thresholds for Models 1, 2 A-D and 3 A-C were determined empirically. The thresholds for Model 4 were determined during parameter estimation.

| MODEL | CELL TYPE | CELL BEHAVIOR | CONDITION |
|---|---|---|---|
| **MODEL 1** | cambium | growth to division threshold | cell area < 10x base cell area |
| | | division | cell area > base area |
| | | | AND [$PXY_{active}$*] > k6 |
| | | differentiation to xylem | cell area > base area |
| | | | AND [$PXY_{active}$ *] < i5 |
| | xylem | growth to max. size | cell area < 10x base cell area |
| | phloem | growth to max. Size | cell area < 10x base cell area |
| **MODEL 2** | cambium | growth to division threshold | cell area < 10x base cell area |
| | | division | cell area > division threshold |
| | | | AND k7 > [$PXY_{active}$ *] > k6 |
| | | differentiation to xylem | cell area > 1.2x rel_cell_div_threshold x base cell area |
| | | | AND [$PXY_{active}$ *] < i5 |
| | | *else if:* differentiation to phloem | cell area > 2x rel_cell_div_threshold x base cell area |
| | xylem | growth to max. size | cell area < 10x base cell area |
| | phloem | growth to max. size | cell area < 10x base cell area |
| **MODEL 3** | cambium | growth to division threshold | cell area < k10x base cell area |
| | | division | cell area > base cell area |
| | | | AND [DF*] > k6 |
| | | *else if:* differentiation to xylem | cell area > 1.9x rel_cell_div_threshold x base cell area |
| | | | AND [PXY*] > k12 |
| | | *else if:* differentiation to phloem | [DF*] > k5 |
| | xylem | growth to max. size | cell area < 1x base cell area |
| | phloem parenchyma | growth to max. size | cell area < 3x base cell area |
| | | division | cell area > 1x base cell area |
| | | | AND [DF*] > k0 |
| | | differentiation to phloem poles | cell area > 1.5x base cell area |
| | | | AND [PF*] < k4 |
| | | | AND cell is not at the tissue boundary |
| | phloem pole | growth to max. size | cell area < 3x base cell area |

| | | | | |
|---|---|---|---|---|
| **MODEL 4** | cambium | growth to division threshold | | cell area < k10 x base cell area |
| | | division | | cell area > k16 x base cell area |
| | | | | AND [DF*] > k6 |
| | | *else if:* | differentiation to xylem | cell area > k11 x rel_cell_div_threshold x base cell area |
| | | | | AND [PXY*] > k12 |
| | | *else if:* | differentiation to phloem | [DF*] > k5 |
| | xylem | growth to max. size | | cell area < k1 x base cell area |
| | phloem parenchyma | growth to max. Size | | cell area < k7 x base cell area |
| | | division | | cell area > 1x base cell area |
| | | | | AND [DF*] > k0 |
| | | differentiation to phloem poles | | cell area > k8 x base cell area |
| | | | | AND [PF*] < k4 |
| | | | | AND cell is not at the tissue boundary |
| | phloem pole | growth to max. size | | cell area < k9 x base cell area |
