## Supplementary File 2 for "Computational modelling of cambium activity provides a regulatory framework for simulating radial plant growth"

### Overview of model parameters and the parameterizations of Model 4.

| PARAMETER | DESCRIPTION | SET1 | SET2 | SET3 | SET4 | SET5 |
|---|---|---|---|---|---|---|
| **K0** | chemical 6 limit above which parenchyma cells can divide | 9.73E-05 | 0.000129 | 0.000122 | 0.000287 | 0.000262 |
| **K1** | xylem maximal cell size | 2.92509 | 1.31994 | 0.865114 | 1.83943 | 0.952836 |
| **K2** | cell growth rate | 5 | 5 | 5 | 5 | 5 |
| **K3** | used to scan the different cell wall stability values of the xylem stiffness regime | NA | NA | NA | NA | NA |
| **K4** | chemical 5 limit below which parenchyma can convert to phloem poles | 2.57E-06 | 1.44E-06 | 1.59E-06 | 2.77E-06 | 2.70E-06 |
| **K5** | chemical 6 limit above which cambium cells can turn into parenchyma | 0.037693 | 0.033071 | 0.045788 | 0.042757 | 0.068131 |
| **K6** | chemical 6 limit above which cambium cells can divide | 5.62E-05 | 0.000182 | 0.000133 | 4.01E-05 | 0.000137 |
| **K7** | phloem parenchyma maximal cell size | 4.8161 | 8.83849 | 5.07855 | 7.4622 | 3.21772 |
| **K8** | size above which parenchyma is converted to phloem poles | 1.9784 | 2.12481 | 2.67309 | 0.688828 | 1.06987 |
| **K9** | phloem pole maximal cell size | 6.15033 | 3.19581 | 4.50996 | 6.28603 | 6.98288 |
| **K10** | cambium maximal cell size | 5.24617 | 5.2554 | 4.55707 | 4.57474 | 2.44148 |
| **K11** | cambium cell size limit above which it can convert to xylem | 0.896592 | 0.818805 | 1.10959 | 1.00441 | 1.15239 |
| **K12** | chemical 1 limit above which cambium cell can convert into xylem | 9.33447 | 9.57172 | 3.24409 | 4.2511 | 3.0528 |
| **K13** | inhibition constant of how much chemical 4 supresses PXY expression | 110.908 | 223.966 | 243.556 | 39.2475 | 127.374 |
| **K14** | rate of how much PXY stimulates the production of DF (pxy mutant = 0) | 199.556 | 229.05 | 194.339 | 48.9899 | 276.148 |
| **K15** | cell size above which parenchyma can divide | 0.485773 | 0.466512 | 0.51765 | 0.656142 | 0.488152 |
| **K16** | cell size above which cambium cells can divide | 0.988171 | 0.957868 | 0.906301 | 0.982287 | 1.05698 |
| **K17** | defines the saturation curve for chemical 6 | 0.006903 | 0.016851 | 0.008233 | 0.031205 | 0.011057 |
| **K18** | defines the saturation curve for chemical 6 | 16.2446 | 28.3991 | 13.2511 | 34.7181 | 33.4421 |
| **K19** | cle41 production rate in phloem parenchyma | 0.157406 | 0.125972 | 0.121037 | 0.245297 | 0.257687 |
| **K20** | cle41 production rate in phloem poles | 2.48875 | 2.45376 | 1.76523 | 2.88386 | 0.935926 |
