## Appendix 1 for "Computational modelling of cambium activity provides a regulatory framework for simulating radial plant growth"

### Description of the VirtualLeaf simulations

VirtualLeaf allows for models to combine tissue dynamics, cell behavior dynamics and biochemical networks that span between cells. The different modeling scales are simulated in sequence. First, the tissue dynamics are simulated using Monte Carlo simulations until a stable energy is reached. Only then are the biological rules applied, with cell division occurring last in order to prevent new cells from interfering with the simulations.

For a detailed description of the simulation process see Merks et al. (2011, 2013) and Antonovici et al. (2022). Here, we include a brief overview of tissue simulations in VirtualLeaf and outline the changes we made for Model 4 as well as the biological rules of the different cambium model versions. The base VirtualLeaf source code is available for download from https://github.com/rmerks/VirtualLeaf2021. The custom version of VirtualLeaf that we built for this analysis as well as the models described in this paper are available at https://github.com/thomasgreb/Lebovka-et-al_cambium-models.

### Tissue simulations

The tissue dynamics are simulated using Monte Carlo simulation dynamics. Briefly, VirtualLeaf attempts to move all nodes of the model in a random order. A Hamiltonian operator is used to assess the energy of the system at both the old and the new position of the node. The movement of nodes is accepted if it minimizes the energy of the system. This operator considers both the cells' compression and the resistance of the cell wall elements to being stretched or compressed (Merks et al. 2011):

$$H = \lambda_A \sum_i (a(i) - A_T(i))^2 - \lambda_M \sum_j (l(j) - L_T(j))^2$$

with $\lambda_A$ as the cell's resistance to compression or expansion, $\lambda_M$ the spring constant for the cell wall elements, $A_T$ and $L_T$ are the cell's target area and the cell wall's target length, respectively, with $a(i)$ representing the current cell area and $l(j)$ the current wall length. For Models 1-3C the standard implementation of the Hamiltonian operator was used.

Cellular growth is implemented in VirtualLeaf as an increase in the cells' target areas. Until the maximal cell size is reached, a cell's target area $A_T(i)$ is increased by a fixed amount in each simulation step. This results in increasing the contribution of the area compression to the Hamiltonian operator

For Model 4 the calculation of the Hamiltonian was refined to include a more detailed definition of the second term for the calculation of the cell wall component of the system's energetic state:

$$\lambda_M \sum \frac{(\lambda_{L1} + \lambda_{L2})}{2} (l(j) - L_T(j))^2$$

Here, $\lambda_{L1,2}$ are cell specific spring constants for the cells that share each specific wall element $j$. Specifically, $\lambda_{L1}$ and $\lambda_{L2}$ are relative contributions to the stiffness of the joint cell wall, where each contribution represents the half of the cell wall secreted by that particular cell. To make the cell wall module compatible with earlier VirtualLeaf models, the default value for $\lambda_L$ is set to "1" such that the expanded calculations result in a multiplication by "1" and do not affect the calculations of the Hamiltonian. The changes to the code in our custom version of VirtualLeaf are marked by a comment "Lebovka et al" at the respective lines of code.

Take the cellular layout of figure 1 as exemplary situation, where node 5 is being moved. During the calculations of the cell wall elements, there are three walls to consider: between nodes 5 and 6, between 5 and 7, and between 5 and 4. As indicated by the arrows, each cell wall will be considered twice during the calculations for the move of node 5: The cell wall between node 4 and node 5 will be called once for cell 1 and once for cell 3, taking into account the specific cell wall thickness specific for each cell.

Altogether, this allows a cell type specific representation of the stiffness of the cell wall elements and therefore a more realistic representation of tissue structure such as an increased cell wall thickness and stability of xylem cells.

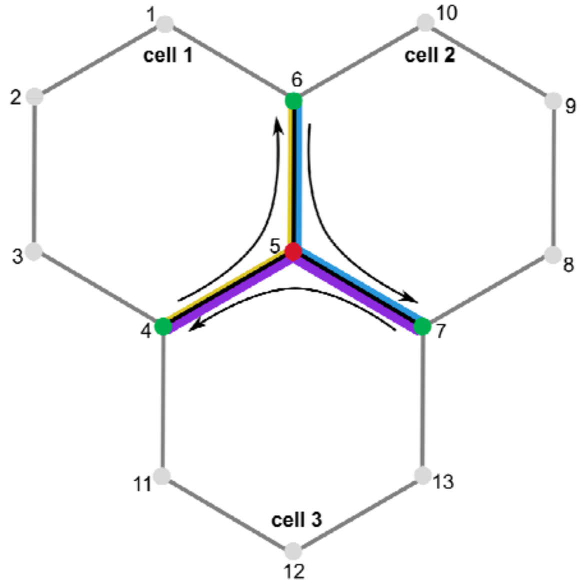


**Appendix-figure 1 - Cell wall calculations during node movement.** Node 5 is moved to a new position. During calculations the change in wall elements between nodes 5 and 6, 5 and 7 as well as 5 and 4 is considered. The cell specific stability of the wall elements is indicated by the thickness of the colored lines – yellow for cell 1, blue for cell 2 and purple for cell 3.

As all the nodes are moved in a random order, this may cause some variation on the tissue layout between simulations. As a consequence, the application of cell behavior rules can vary as well between simulation runs, e.g. as cells divide over the longest axis and not in a predetermined direction. To account for these variations between simulations, we simulated each model, each parameter set and each thickness regime at least ten times.

### Cambium Models

Models in VirtualLeaf comprise four different files: (1) the project file, (2) the model header file, (3) the C++ file containing the model algorithm and (4) the tissue layout. We provide these four files for the cambium models in the GitHub repository linked above. The model will further need to be included in the Model.pro file as a subdirectory by including the line “Model_folder \” as one of the entries below “SUBDIR = \”.

#### 1. Model.pro

This is a C++ project file containing the configuration settings and pathways for the necessary directories.

#### 2. Model.h

This is a C++ header file containing a line with the following structure: “virtual QString DefaultLeafML(void) {return QString("hypo7.xml");}”. The line indicates where VitualLeaf should search for an xml file that describes the structure of the tissue template (called “leaf”) used for the model to run upon. In this particular example, the name of the xml template is “hypo7.xml”. VirtualLeaf will go to the folder in which you installed the software and will look for this file in the subfolder “../data/leaves”. In our case, a Windows machine was used. Therefore, the full path looked like this:

"C:\VirtualLeaf2021-main\data\leaves" and this folder contained a file "hypo7.xml". Please note that paths will be different depending on the operating system being used.

**2. Model.cpp**

A C++ file containing the model algorithm to reproduce the output described in this study. Each model contains specific rules for cell behavior and biochemical equations specific to the cell types defined in the leaf.xml file. The cell behavior rules are listed in the sections *OnDivide* and *CellHousekeeping* while the biochemical model is listed in the section *CellDynamics*. Cell-to-cell transport is considered in the section *CelltoCellTransport* with reactions at cell walls having their specific section *WallDynamics*, though the latter was not used in any of the Cambium models.

**2.1 Cell behavior models**

All cells in the cambium models follow specific behavioral rules governing cell growth, cell division and cell differentiation (Tab. S1). Generally, cells grow until a maximal size is reached, unless other behavior rules are triggered. Cell division and differentiation require not only a minimal cell size but also additional conditions regarding chemical concentrations. Unless otherwise specified, all cell behavior rules are applied as long as the specific conditions are met.

**2.2 Biochemical Model**

**Model 1 & 2**

In cambium* and xylem* cells, CLE41 dynamics are a combination of the diffusion of CLE41*, the binding to PXY* and the degradation of CLE41*:

$$\frac{d}{dt}[CLE41^*] = \ diffusion_{CLE41} - [PXY^*] \cdot [CLE41^*] - degradation_{CLE41} \cdot [CLE41^*]$$

In phloem* cells, there is an additional term in the equation describing the production of CLE41*:

$$\frac{d}{dt}[CLE41^*] = \ diffusion_{CLE41} + production_{CLE41} - [PXY^*] \cdot [CLE41^*]$$

$$-degradation_{CLE41} \cdot [CLE41^*]$$

PXY* is produced in cambium* cells and negatively regulated by bound PXY*:

$$\frac{d}{dt}[PXY^*] = \frac{production_{PXY}}{(1 + suppress\ rate \cdot [PXY^*_{active}])} - [PXY^*] \cdot [CLE41^*]$$

$$-degradation_{PXY} \cdot [PXY^*]$$

In the other cell types* in turn, free PXY* is governed by CLE41* binding to PXY* as well as the degradation of the receptor:

$$\frac{d}{dt}[PXY^*] = -[PXY^*] \cdot [CLE41^*] - degradation_{PXY} \cdot [PXY^*]$$

The ODE describing the dynamics of bound PXY* is identical for all cell types*. Here, bound PXY* is produced by the association of CLE41* and PXY and later degraded:

$$\frac{d}{dt}[PXY^*_{active}] = [PXY^*] \cdot [CLE41^*] - degradation_{PXY^*_{active}} \cdot [PXY^*_{active}]$$

**Model 2B**

In Model 2B CLE41* is also produced in xylem cells*, such that the ODE now reads as follows:

$$\frac{d}{dt}[CLE41^*] = \ diffusion_{CLE41} + production_{CLE41} - [PXY^*] \cdot [CLE41^*]$$

$$-degradation_{CLE41} \cdot [CLE41^*]$$

**Model 2C & D**

In Model 2C and D the production of PXY* in cambium cells is eliminated (C) or strongly reduced (D). As such, the ODE for PXY* in Model 2D is now:

$$\frac{d}{dt}[PXY^*] = \frac{0.1 \ \cdot production_{PXY}}{(1 + suppress\ rate \cdot [PXY^*_{active}])} - [PXY^*] \cdot [CLE41^*]$$

$$-degradation_{PXY} \cdot [PXY^*]$$

For Model 2C the production term is set to "0", fully eliminating PXY* production in cambium cells*.

**Models 3 & 4**

In Models 3 and 4 we expanded the biochemical network to include additional chemicals suppressing PXY expression (RP*), a dedicated division factor as well as phloem derived factors promoting the division factor and suppressing phloem pole formation ($PF_{div}$* and $PF_{pole}$*, respectively). While the ODEs for CLE41*, free PXY* and bound PXY* remain mostly unchanged, we refined the ODE for PXY* to make the production of PXY* independent of $PXY_{active}$*:

$$\frac{d}{dt}[PXY^*] = \frac{production_{PXY}}{\left(1 + suppress\ rate \cdot \left[RP^*\right]\right)} - [PXY^*] \cdot [CLE41^*]$$

$$-degradation_{PXY} \cdot [PXY^*]$$

We also set the production rates of CLE41* to be higher in phloem poles* than in phloem parenchyma*.

The factor suppressing PXY expression (RP*) diffuses and is degrade throughout the tissue but is only produced in phloem cells. We therefore get in the following equation for phloem cells:

$$\frac{d}{dt}[RP^{*}] = production_{RP} + diffusion_{RP} - degradation_{RP} \cdot [RP^{*}]$$

In all other cell types, this ODE is simplified to include only the diffusion and degradation of RP*.

For the second phloem-derived factor, PF*, two chemicals were defined in the biochemical model on account of the different functions in the model reminiscent of different signaling components in planta: promoting the production of the division chemical reminiscent of the PEAR transcription factors ($PF_{div}$*) and suppressing phloem pole formation reminiscent of the CLE45/ RPK2 signaling module ($PF_{pole}$*). The respective ODEs for both $PF_{div}$* and $PF_{pole}$* in phloem poles* are therefore:

$$\frac{d}{dt}[PF_{div}^{*}] = production_{PF} + diffusion_{PF} - degradation_{PF} \cdot [PF_{div}^{*}]$$

$$\frac{d}{dt}\left[PF_{pole}^{*}\right] = production_{PF} + diffusion_{PF} - degradation_{PF} \cdot \left[PF_{pole}^{*}\right]$$

In all other cell types, these ODE are simplified to include only the diffusion and degradation of $PF_{div}$* and $PF_{pole}$*.

Last, we included a factor promoting the division of cambium* and phloem parenchyma* cells (DF*). Generally, the division chemical DF* is degraded in tissues:

$$\frac{d}{dt}[DF^{*}] = diffusion_{DF} - degradation_{DF} \cdot [DF^{*}]$$

Only, in phloem parenchyma* and cambium* cells this chemical is also produced:

$$\frac{d}{dt}[DF^*] = \frac{production_{DF} \cdot ([PF^*] + 100 * [PXY^*_{active}])}{K + [PF^*] + 100 * [PXY^*_{active}]} + diffusion_{DF}$$

$$-degradation_{DF} \cdot [DF^*]$$

**Model 3B**

In Model 3B CLE41* is also produced in xylem cells*, such that the ODE now reads as follows:

$$\frac{d}{dt}[CLE41^*] = diffusion_{CLE41} + production_{CLE41} - [PXY^*] \cdot [CLE41^*]$$

$$-degradation_{CLE41} \cdot [CLE41^*]$$

**Model 3C**

In Model 3C, the implementation of the *pxy* mutant was two-fold, as we needed PXY* in the model for the positional information during xylem cell* differentiation. First, the production of $PXY^*_{active}$ was set to zero. And second, the DF* production only depended on DF*:

$$\frac{d}{dt}[DF^*] = \frac{production_{DF} \cdot ([PF^*])}{K + [PF^*]} + diffusion_{DF} - degradation_{DF} \cdot [DF^*]$$

**Diffusion**

Generally, we defined the diffusion flux $phi$ according to Fick's law, i.e. based on the concentrations of neighboring cells and the length of the shared cell wall element

$$phi = length_{wall\ element} \cdot (concentration_{cell\ 2} - concentration_{cell\ 1}),$$

so that the change in cell 1 is equal to $phi$ and the change in cell 2 is equal to $-phi$.

In Model 1 only CLE41 diffuses between cells with no restrictions regarding to cell types. In Models 3 and 4 we also considered the diffusion of RP*, $PF_{div}$*, $PF_{pole}$* and DF*, all of which were calculated according to the equation above and without restrictions regarding to cell types.

#### 3. Leaf.xml

A file containing the description of a tissue template as described before (Merks *et al.* 2011). The software uses this file to construct a tissue template and to run a given model.

In order to run or modify a provided model, follow the following instructions.

**a.** Create a new model with the desired name (e.g. “my_cool_model”) as described (Merks *et al.* 2011).

**b.** After a new model was created, there should be a folder “../src/Models/my_cool_model” in your VirtualLeaf folder. In our case, the full path looked like this: “C:\VirtualLeaf2021-main\src\Models\ my_cool_model”.

**c.** In your “../src/Models/my_cool_model” folder locate “my_cool_model.h” and “my_cool_model.cpp” files. Using a text editor replace the content of those files by the content of the respective files from the model you are interested in (files provided in this paper are called “Model1.h” and “Model1.cpp”). Please note that you should only replace the content of the files and not the files themselves. After you have completed this step, your files should still be named “my_cool_model.h” and “my_cool_model.cpp”.

**d.** Open the files “my_cool_model.h” and “my_cool_model.cpp” using a text editor and replace every instance of “Model1” by “my_cool_model” in the text. Save the changes.

**e.** Locate the "../data/leaves" folder and add the provided xml file defining the tissue template (in our case, the tissue template is called "hypo7.xml'). The resulting full path to the file had the following structure in our case: "C:\VirtualLeaf2021-Main\data\leaves\hypo7.xml".

**f.** Compile the model as described (Merks *et al.* 2011, Antonovici *et al.* 2022). Please note that each time you introduce changes into the code, you should recompile the model and re-start VirtualLeaf.

**g.** Now you can run VirtualLeaf. Go to the "../bin' folder and run the "VirtualLeaf" file. In our case the full path looked like this: "C:\VirtualLeaf2021-main\bin\VirtualLeaf ".

The new model will appear under the "Models" section with the corresponding name. Please note that the name of the model that will be shown is not the same as "my_cool_model". Instead, it will show whichever name was indicated in the "my_cool_model.cpp" file in this line: // specify the name of your model here; return QString( "Model 1 - pxy only" )". In this case, there will be a new model called "Model 1 - pxy only" in the VirtualLeaf folder under the "Models" section.
